# Cancer-associated Fibroblast Spatial Heterogeneity and *EMILIN1* Expression in Cancer Stroma Modulate TGF-β Activity and CD8^+^ T-Cell Infiltration in Breast Cancer

**DOI:** 10.1101/2023.09.12.557312

**Authors:** Chikako Honda, Sasagu Kurozumi, Takaaki Fujii, Didier Pourquier, Lakhdar Khellaf, Florence Boissiere, Jun Horiguchi, Tetsunari Oyama, Ken Shirabe, Jacques Colinge, Takehiko Yokobori, Andrei Turtoi

**Affiliations:** Department of General Surgical Science, Gunma University Graduate School of Medicine, Maebashi, Gunma, Japan; Department of Breast Surgery, International University of Health and Welfare, Chiba, Japan; Institut régional du Cancer de Montpellier (ICM)-Val d’Aurelle, Montpellier, France; Tumor Microenvironment and Resistance to Treatment Lab, INSERM U1194, Montpellier, France; Université de Montpellier, Montpellier, France; Department of Pathology and Diagnostics, Gunma University Graduate School of Medicine, Maebashi, Gunma, Japan; Cancer Bioinformatics and Systems Biology Team, INSERM U1194, Montpellier, France; Division of Integrated Oncology Research, Gunma University, Initiative for Advanced Research (GIAR), Maebashi, Gunma, Japan

**Author notes:** Co-corresponding authors: Andrei Turtoi, PhD; Takehiko Yokobori, MD, PhD; Jacques Colinge, PhD. contributed equally (Jacques Colinge, Takehiko Yokobori, Andrei Turtoi).

## Abstract

The tumor microenvironment (TME) and its multifaceted interactions with cancer cells are major targets for cancer treatment. Single-cell technologies have brought major insights into the TME, but the resulting complexity frequently precludes conclusions on function. Therefore, we combined single-cell RNA sequencing and spatial transcriptomic data to explore the relationship between different cancer-associated fibroblast (CAF) populations and immune cell exclusion in breast tumors. Our data show for the first time the degree of spatial organization of different CAF populations in breast cancer. We found that IL-iCAFs, Detox-iCAFs, and IFNγ-iCAFs tended to cluster together, while Wound-myCAFs, TGFβ-myCAFs, and ECM-myCAFs formed another group that overlapped with elevated TGF-β signaling. Differential gene expression analysis of areas with CD8^+^ T-cell infiltration/exclusion within the TGF-β signaling-rich zones identified elastin microfibrillar interface protein 1 (*EMILIN1*) as a top modulated gene. *EMILIN1*, a TGF-β inhibitor, was upregulated in IFNγ-iCAFs directly modulating TGFβ immunosuppressive function. Histological analysis of 74 breast cancer samples confirmed that high EMILIN-1 expression in the tumor margins was related to high CD8^+^ T-cell infiltration, consistent with our spatial gene expression analysis. High EMILIN-1 expression was also associated with better prognosis of patients with breast cancer, underscoring its functional significance for the recruitment of cytotoxic T cells into the tumor area. In conclusion, our data show that correlating TGF-β signaling to a CAF subpopulation is not enough because proteins with TGF-β-modulating activity originating from other CAF subpopulations can alter its activity. Therefore, therapeutic targeting should remain focused on biological processes rather than on specific CAF subtypes.

## INTRODUCTION

Breast cancer (BC) is the second most frequent cause of cancer death in women worldwide^1^. Molecular and histological classifications of BC have significantly improved its clinical management. Today, histopathological assessment of needle biopsies for morphological type and histological grade determination are complemented by the assessment of estrogen receptor (ER), progesterone receptor (PgR), human epidermal growth factor receptor type 2 (HER2) status, and Ki-67 proliferative index^2^. Therefore, BC are classified into three types: (a) hormone receptor (ER and/or PgR)-positive and HER2-negative, (b) HER2-positive, and (c) hormone receptor-and HER2-negative (triple negative breast cancer, TNBC)^2^. Based on this classification, drug treatment regimens for early invasive BC include endocrine therapy (a) and chemotherapy and anti-HER2 agents (b), alone or in combination^3^. No specific treatment is available for TNBC (c), and this represents a tremendous clinical challenge^3,4^. Indeed, no druggable vulnerability has been identified in TNBC cells to date, precluding their direct targeting^4,5^.

A complementary approach to cancer cell targeting is to target the environment in which they reside^6^ and that is called tumor microenvironment (TME). The TME consists of a cellular part (stroma) and a supportive extracellular matrix (ECM) with specific physical and chemical properties. The stroma is primarily composed of endothelial, immune and fibrotic components. All three have attracted considerable attention for novel drug development in solid tumors. Indeed, in some cancers, targeting endothelial and immune cells is more effective than killing cancer cells (e.g. melanoma^7,8^ and hepatocellular^9^ carcinoma). This demonstrates the potential of TME-directed therapies, possibly in combination with molecules against cancer cells. Despite these encouraging results, BC (like many other solid tumors) has not really benefited from TME targeting yet. Clinical trials produced rather mitigated results. For example, in BC, endothelial cells have been mainly targeted with anti-angiogenic drugs (e.g. anti-vascular endothelial growth factor (VEGF) antibodies), alone or in combination with chemotherapy^10,11^. Unfortunately, the survival benefit for patients with BC was minimal, and several potential resistance mechanisms were described^12^. The immune component of BC has been mainly targeted using novel monoclonal antibodies against immune checkpoint proteins (e.g. programmed cell-death protein 1 (PD-1) and its ligand PD-L1) with the aim of restoring the anti-tumor immunity. The immune checkpoint inhibitors (ICI) pembrolizumab (anti-PD-1) and atezolizumab (anti-PD-L1) have been clinically tested in patients with metastatic TNBC with heterogeneous results^13,14^. A good clinical response is achieved in a small subpopulation of patients, and no clear biomarker exists to predict which patients will respond. Patients with BC characterized by high mutational burden or with immunologically inflamed tumors (with high proportion of CD8^+^ T cells in the tumor center) might respond to ICIs^15,16^. Indeed, ICIs that activate cytotoxic T cells against tumors are now considered an important therapeutic tool^17,18^. Lastly, clinical trials on cancer-associated fibroblasts (CAFs) and the targeting of the tumor fibrotic component have not brought any conclusive results in BC, leaving this area unexplored. Despite the plethora of experimental data on CAF significance in BC progression^19^, very few targetable molecules have been identified in CAFs. Additionally, recent data on CAF subpopulations in BC^20^ raise the question of whether specific subtypes should be targeted.

However, we believe that successful TME targeting should be process-oriented and not cell-oriented. Indeed, immune exclusion and angiogenesis are promoted and regulated by the concerted action of several stromal cell types, and therefore disrupting a specific cell population is unlikely to abolish such processes entirely. Conversely, it might be more relevant to target the potentially trans-cellular molecular network at the heart of a crucial tumor process. However, knowledge on this area is still limited. Particularly, it is crucial to understand how different stromal cells molecularly engage to support tumor-promoting programs.

Recent advances in spatial OMICS technologies have been a true game changer for characterizing the TME and tumor heterogeneity^21^. They have allowed, for the first time, to link the spatial occurrence of different TME cell subtypes and of cancer cells with enhanced proliferative or therapy-resistance features^22^. In the present study, we investigated the spatial distribution of CAF subpopulations in BC and their relationship with infiltrating cytotoxic T cells.

## MATERIALS AND METHODS

### Patient Material

Four patients with invasive ductal BC who underwent surgical resection at Gunma University Hospital (Gunma, Japan) in 2020-2021 were enrolled for the Visium Spatial Gene Expression experiments (clinical data are in **Table S1**). For immunohistochemical staining (validation study), 75 patients with invasive BC who underwent breast-conserving surgery or modified total mastectomy at Gunma University Hospital (Gunma, Japan) in 2020-2021 were enrolled (**Table S2**). Men with BC were not included in the study. None of the patients received neoadjuvant treatment. Their median age was 60 years (range, 35-82 years). Pathological tumor size, nodal status and lymphovascular invasion were determined using the pathological records. The present study was approved by the Gunma University Hospital Institutional Review Board (reference no. HS2021-071) and was conducted according to the tenets of the Declaration of Helsinki. All patients gave their consent via the opt-out system.

### Tissue Optimization

Tissue optimization was performed following the 10x Genomics Visium Spatial Tissue Optimization Reagents Kits User Guide (CG000238, 10x Genomics) to optimize the permeabilization time for the subsequent gene expression profiling. BC tissue cryosections (10 μm-thick) were placed on a Visium Spatial Tissue Optimization Slide (10x Genomics). Different permeabilization times were tested with different tissue sections on the slide with poly(dT) primers to capture the mRNA. After the permeabilization and the mRNA capture steps, reverse transcription followed by addition of fluorescently labeled oligonucleotides to the cDNA allowed detecting the resulting cDNAs as fluorescence signals. Hematoxylin and eosin (H-E) staining and the fluorescence signals were imaged with a BZ-X800 microscope (Keyence). The optimal permeabilization time was the incubation time that gave the strongest fluorescence signal.

### Gene expression analysis library preparation

Spatial gene expression analysis was done with the Visium Spatial Gene Expression Reagent Kit (10x Genomics) following the manufacturer’s user guide (CG000239, 10x Genomics). BC tissue cryosections (10 μm-thick) were placed on a Visium Spatial Gene Expression Slide (10x Genomics). Images of H-E-stained sections were taken with a BZ-X800 microscope (Keyence). After tissue permeabilization for the optimal time (see above), mRNA capture with the poly(dT) probes in the slide and reverse transcription resulted in the construction of the full-length cDNA. After second strand synthesis and denaturation, cDNAs were amplified in a Veriti 96-Well Thermal Cycler (Thermo Fisher Scientific) and quantified with a LabChip GX Touch HT Nucleic Acid Analyzer (PerkinElmer) to ensure that sufficient cDNA amounts were generated for the library construction. Enzymatic fragmentation and size selection with the SPRIselect reagent (Beckman Coulter) were used to optimize the cDNA fragment size for sequencing. Then, sample index PCR allowed preparing sequence-ready libraries. The final library quantification was done with LabChip GX.

### Sequencing

The MGIEasy Universal Library Conversion Kit (MGI Tech) was used to convert the libraries to DNBSEQ-compatible libraries. Sequencing was done by DNBSEQ-G400 (MGI Tech) with a DNBSEQ-G400RS High-throughput Sequencing Set (App-A FCL PE100) following the manufacturer’s instructions. The resulting read lengths were as follows: Read1-28 bp and Read2 -100bp.

### Bioinformatics

The raw fastq files were processed with the SpaceRanger software 1.0 (10x Genomics) using the human genome reference set GRCh38-3.0.0 and default parameters. Data obtained from our four BC samples were complemented with published data^23^ retrieved from GEO (reference GSE176078). For our four samples, tissue areas were defined using the Seurat clustering default algorithm (functions FindNeighbors and FindClusters). The cluster number was adjusted to the maximum value where distinct, cluster-specific gene expression patterns were detected with the Seurat differential search tool (function FindAllMarkers). These tissue areas were named by referring to the original areas defined by a pathologist. For the publicly available datasets^23^, the original area definitions were used. Of note, these tumor areas played no role in the analysis, and they were defined only for reference and descriptive purposes.

The gene spatial expression analysis mainly relied on our library BulkSignalR^24^ and project-specific R scripts. Count matrices were filtered for non-expressed genes by imposing a minimum read count of 1 in at least 1% of the Visium spots. Subsequently, normalization was achieved by total count. Cell population-specific gene signatures were retrieved from sequence data for BC general cell populations^23^ and for CAFs^25^. In all cases, the top 20 genes reported for each population were used. The spatial abundance of each cell population was estimated by applying BisqueRNA^26^ to these gene signatures due to the bulk nature of Visium spatial data. A first scoring of cycling cancer cells was obtained using BisqueRNA scores for the Cycling population^23^. An alternative score was provided by scoring a gene signature available from Seurat (cc.genes.updated.2019$s.genes and cc.genes.updated.2019$g2m.genes). In this case, scoring was done using the BulkSignalR function scoreSignatures. The various plots reporting the localization of cell types or cycling cells were generated using BulkSignalR standard functions.

Several biological processes were scored using fast Gene Set Enrichment Analysis (fGSEA)^27^ spatial transcriptomic features. TGF-β signaling was scored with the BulkSignalR scoreSignatures function to generate dendrograms to relate this process to CAF subpopulations. TGF-β signaling genes were obtained from MSSigDB (c5.bp.v7.0.symbols.gmt.txt, GO_TRANSFORMING_GROWTH_FACTOR_BETA_RECEPTOR_SIGNALING_PATHWAY). Spatial co-localization between cell populations or between TGF-β and cell populations was determined using a Pearson correlation-based distance matrix (distance = 1 – correlation). Correlations within one sample were computed over the whole set of spots.

The differential gene expression analysis to compare CD8^+^ T cell-rich *versus* -poor areas with the top TGF-β signaling tumor areas was performed with edgeR (PMID: 22287627) and the following parameters: maximum false discovery rate of 5%, minimum fold-change of 1.5, and normalized read count >1 in at least 25% of spots. For one sample (tumor 114223F), this last threshold was decreased to 20%.

### Immunohistochemistry

Paraffin-embedded BC specimens (n=75; 10 luminal A, 10 luminal B, 20 luminal HER2, 15 HER2, and 20 TNBC) were cut into 4 μm-thick sections and mounted on glass slides. All sections were incubated at 60ºC for 60 min, deparaffinized in xylene, rehydrated, and incubated with fresh 0.3% hydrogen peroxide in 100% methanol at room temperature for 30□min to block endogenous peroxidase activity. After rehydration through a graded series of ethanol solutions, antigen retrieval was performed using an Immunosaver (Nishin EM, Tokyo, Japan) at 98º C-100°C for 30□min. Sections were passively cooled to room temperature and then incubated in Protein Block Serum-Free Reagent (Agilent (Dako), Santa Clara, CA, USA) for 30 min. This was followed by incubation with an anti-EMILIN-1 rabbit polyclonal antibody (x400, HPA002822; Sigma Aldrich, Saint Louis, MO, USA) in Dako REAL Antibody Diluent at 4°C for 24 h. According to the manufacturer’s instructions, EMILIN-1 staining was visualized as a red color using the Histofine Simple Stain AP (Multi) Kit (Nichirei, Tokyo, Japan) and the FastRed II reagent (Nichirei, Tokyo, Japan). Then, sections were boiled in a microwave oven for 10 min to inactivate the antibodies and enzyme activity. Next, they were incubated with an anti-CD8 rabbit polyclonal antibody (x500, ab4055; Abcam, Cambridge, UK) in Dako REAL Antibody Diluent at 4°C for 24 h. CD8 staining was visualized as a brown color using the Histofine Simple Stain MAX-PO (Multi) Kit (Nichirei, Tokyo, Japan) and DAB substrate. Sections were lightly counterstained with hematoxylin and mounted. Negative controls were incubated without the primary antibody, and no staining was detected.

EMILIN-1 expression was evaluated as staining intensity and staining ratio in 200x view fields from two tumor margin areas and one center area. Staining intensity was evaluated as 0 (none), 1 (weak), 2 (moderate), and 3 (strong). The ratio of EMILIN-1-stained area to the whole field of view was evaluated as 0 (none), 1 (1%-25%), 2 (26%-50%), 3 (51%-75%), 4 (≧76%). The staining intensity and ratio were multiplied to obtain the EMILIN-1 score (0-12). BC samples with higher EMILIN-1 score in the margin than central area were defined as a high EMILIN-1 group, and the others as low EMILIN-1 group. The total number of CD8^+^ cells was counted in the 200x view fields where the EMILIN-1 score was evaluated.

### Immunofluorescence analysis

Multicolor immunofluorescence staining was performed in tissue sections of BC surgically resected from five patients to detect EMILIN-1, CD8, and TGFBI expression and from seven patients to detect EMILIN-1, CD8, and Ki-67 using the Akoya Biosciences Opal Kit following the manufacturer’s instructions. All patients were selected from the validation group of BC samples. In the first five samples, EMILIN-1 staining (anti-EMILIN-1 rabbit polyclonal antibody: x400, HPA002822, Sigma) was visualized using the Opal 480 Fluorophore; CD8 staining (anti-CD8 rabbit polyclonal antibody: x500, ab4055, Abcam) with the Opal 570 Fluorophore; and TGFBI staining (anti-TGFBI rabbit polyclonal antibody: x400, 10188-1-AP; Proteintech, Rosemont, IL, USA) with the Opal 520 Fluorophore (**Figure 5a**). In the other seven BC samples, EMILIN-1 staining was visualized using the Opal 480 Fluorophore, CD8 staining with the Opal 570 Fluorophore, as above, and Ki-67 staining (anti-Ki67 rabbit monoclonal antibody: x500, #9027; Cell Signaling Technology, Danvers, MA, USA) with the Opal 520 Fluorophore (**Figure 5c**). All sections were lightly counterstained with hematoxylin and examined under an All-in-One BZ-X710 fluorescence microscope (KEYENCE Corporation, Osaka, Japan).

## RESULTS

### The integration of single-cell RNA-seq and spatial transcriptomic data unveils functional heterogeneity across BC samples

We generated spatial transcriptomic data from four untreated invasive BC samples (A1, B1, C1, and D; see **Table S1** for the tumor classification). We also included published data on six BC samples (1160920F [TNBC], 1142243F [TNBC], CID4290 [ER+], CID4535 [ER+], CID4465 [TNBC], CID44971 [TNBC])^23^. To gain a deeper insight into the cellular composition of each BC sample, we decided to assemble a single-cell RNA-seq BC atlas. To this end, we merged data from two recently published studies. The first one characterized all cell populations in 26 BC samples^23^, and the second one characterized >18,000 CAFs from 8 BC samples^25^. To avoid redundancy, we removed the CAF populations from the first study and kept only the CAF populations from the second study. The resulting atlas is featured in **Figure 1a**, and representative cell-specific markers are in **Figure 1b**. Then, we used this BC cell atlas to annotate spatial transcriptomic data for our ten BC samples and to project each cell population on histological sections (**Figures S1-S10**). The annotation relevance was verified relative to the presence of the typical histological structures observed in the H-E stained histological sections (e.g. vasculature, fibrosis, *in situ* versus invasive cancer). An example is shown in **Figure 1c**. This annotation process highlighted that individual cell types compartmentalized differently in different BC samples, suggesting significant functional heterogeneity. To assess this, we performed a spatial gene ontology (GO) analysis using fGSEA. All BC specimens (n=10) showed a significant modulation of >300 biological processes (data not shown) that could be grouped in two to three spatial patterns *per* sample (**Figures 1d & S11**). Among the spatially modulated GO processes, those relating to ECM remodeling and immunity regulation were particularly relevant for studying the CAF-immune cell interactions. Both processes showed a clear tissue compartmentalization, suggesting that such interactions may be enriched in specific BC tissue regions (**Figure 1d**). TGF-β signaling was particularly interesting due to its capacity to suppress tumor immune response^28^. The T-cell and macrophage activation processes showed a distinct compartmentalization, but with a spatial pattern opposite to that of TGF-β signaling. However, some tumor regions were rich in both TGF-β signaling and T-cell activation. This finding was particularly intriguing and required additional analyses.

**Figure 1:**
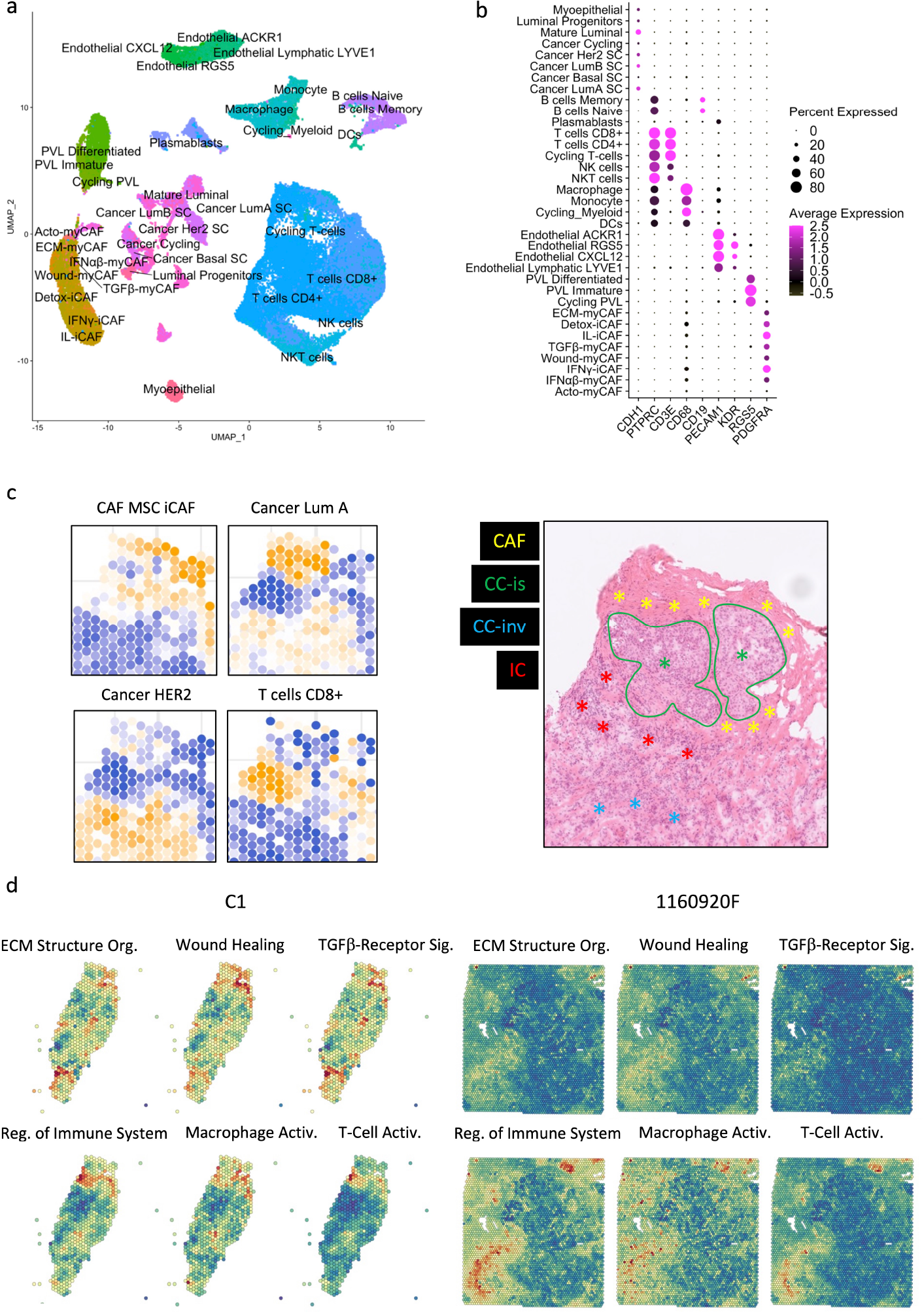
Breast cancer atlas for cellular and functional annotation of spatial single-cell RNA-seq data. (**a**) UMAP plot showing the breast cancer atlas based on two previously published single-cell RNA-seq datasets.^21,23^ (**b**) Verification of the cellular annotation of the newly generated breast cancer atlas using several cell-specific genes. (**c**) Deconvolution of two cell populations in the spatial single-cell RNA-seq data using the breast cancer cell atlas (left panels) and confirmation of the deconvolution by histology analysis (right panel). Different colors denote regions with certain predominant cell types, such as: cancer-associated fibroblasts (CAF), in-situ cancer cells (CC-is), invasive cancer cells (CC-inv) and immune cells (IC). (**d**) Spatial distribution of selected GO processes in BC samples (two representative samples are shown: C1 and 1160920F; other samples are displayed in **Figure S1**). The following GO processes are displayed: ECM Structural Organization, Wound Healing, TGF-β Receptor Signaling, Regulation of Immune System, Macrophage Activation, and T-Cell Activation.

### BC areas with low proliferation potential are characterized by high macrophage and CD8^+^ T-cell infiltration

The spatial transcriptomic data projection on histological sections (**Figures S1-S10**) showed that BC samples were composed of distinct and heterogeneously distributed cancer cell subtypes. However, we observed a high degree of spatial consistency among cycling cancer cells and regions characterized by high-proliferative potential (**Figure S12**, S+G2M score plots). Spatial deconvolution of the immune cell infiltrate indicated that CD8^+^ T cells and macrophages were abundant in regions with low proliferative capacity (**Figure 2a-b**). The correlation analysis confirmed this pattern in 9/10 BC spatial datasets (the only exception was CID4535). As CD8^+^ T cells have a crucial role in tumor growth inhibition, we next determined whether locoregional differences in TGF-β signaling were correlated with the differential presence of CD8^+^ T cells. Indeed, TGF-β is a well-known master regulator of normal and pathologic inflammation. We did not find any significant correlation between TGF-β signaling and CD8^+^ T-cell abundance (data not shown), suggesting a more complex relationship between immune exclusion and TGF-β signaling. CAFs are major TGF-β producers in tumors and also regulate TGF-β activity through the secretion of modulatory proteins^29^. Therefore, we hypothesized that CAF populations and their tissue distribution might explain the link between TGF-β activity and CD8^+^ T-cell exclusion.

**Figure 2:**
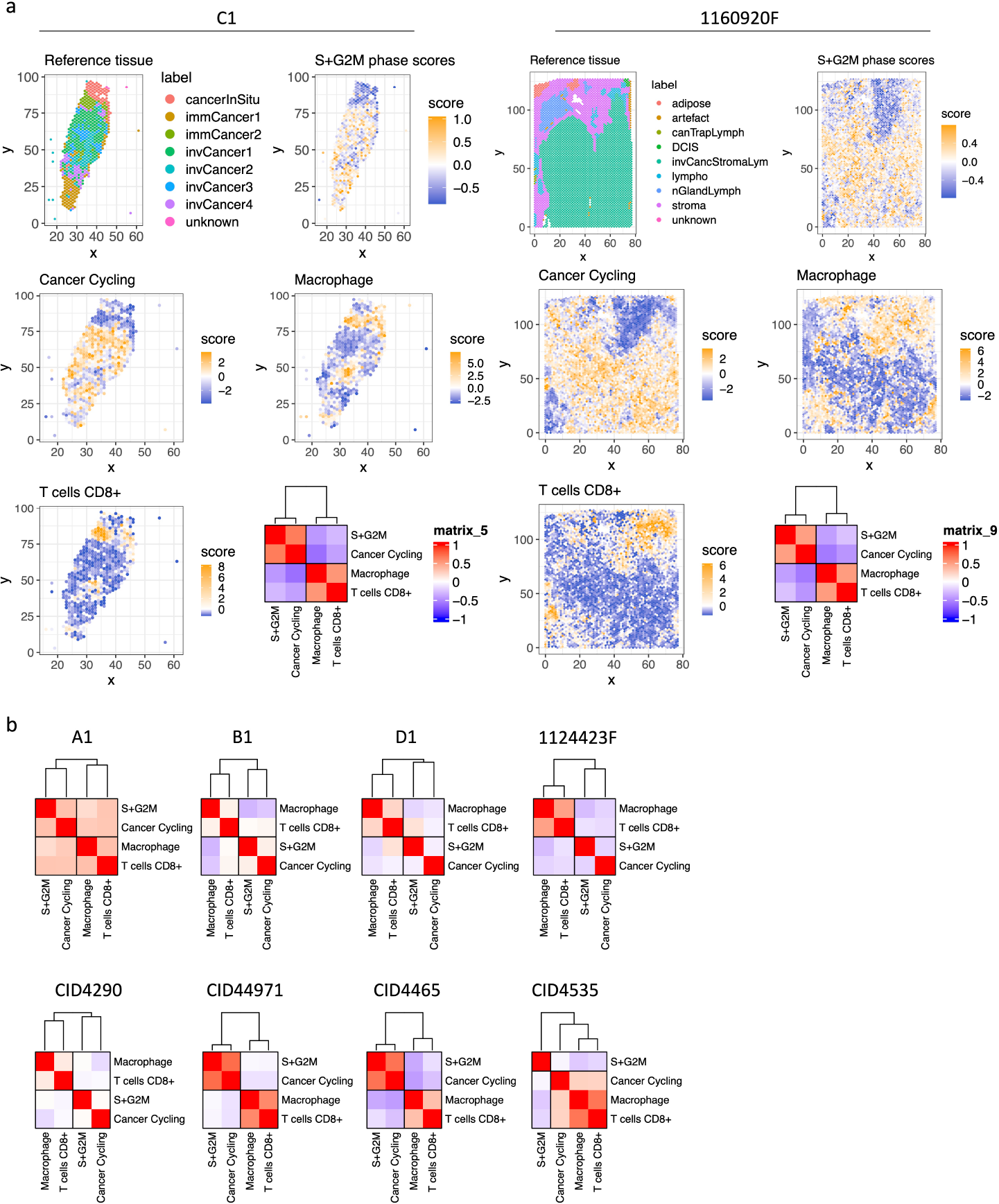
Spatial analysis of proliferating cancer cells and immune infiltrate in breast cancer samples. (**a**) Histological annotation of two representative breast cancer samples (other samples are shown in **Figure S12**), and estimation of highly proliferative regions (S+G2M phases) (higher panels); actively cycling cancer cells and two immune populations (macrophages and CD8^+^ T cells) (middle and lower panels). The heath map shows the correlation analysis for these four populations. (**b**) Correlation analysis for the four selected cell populations in the other eight breast cancer samples.

### ECM-myCAFs, wound-myCAFs and TGFβ-myCAFs are in regions with high TGF-β signaling

Recent single-cell studies^25^ determined that in BC, there are several major CAF subpopulations: ECM-myCAFs, TGFβ-myCAFs, wound-myCAFs, IFNαβ-myCAFs, acto-myCAFs, IFNγ-iCAFs, detox-iCAFs, and IL-iCAFs. These CAF subpopulations are characterized by distinct gene expression profiles that suggest their involvement in specific cancer-relevant biological pathways. An overview of the GO enrichment analysis in each CAF subpopulations is provided in **Figure S13**. No significant acto-myCAF enrichment was observed. This was the smallest CAF population, at the periphery of the CAF cluster in **Figure 1a**. Next, we used single-cell RNA-seq data to spatially map CAF subpopulations in the BC samples. CAF subpopulations showed a rather compartmentalized distribution pattern (**Figure 3a** and **Figures S1-S10**). Overall, their distribution profiles could be classified in two main patterns (**Figure 3a-b**) that included ECM-myCAFs, wound-myCAFs and TGFβ-myCAFs (first pattern) and detox-iCAFs, IL-iCAFs and IFNγ-iCAFs (second pattern). Conversely, IFNαβ-myCAFs frequently grouped separately. We also found that high TGF-β signaling was associated with the first pattern (ECM-myCAFs, wound-myCAFs and TGFβ-myCAFs). Having established a link between these three CAF subpopulations and TGF-β signaling, we wanted to understand its potential effect on the spatial localization of CD8^+^ T cells. Specifically, we asked why CD8^+^ T cells could accumulate in some BC areas that were rich in TGF-β signaling, although this is in contradiction with TGF-β immune suppressor role (**Figure 4a**). We spatially scored TGF-β signaling in each BC sample using a gene set (Materials and Methods), and defined a tumor-specific high TGF-β area that corresponded to the top TGF-β signaling scores. Independently, we defined CD8^+^ T cell-rich and -poor areas in each tumor sample (i.e. the locations with top and bottom quarter CD8^+^ T-cell abundance scores, respectively). By intersecting these areas, we compared gene expression in CD8^+^ T cell-rich *versus* -poor areas within high TGF-β signaling locations (**Figure 4a**). Differential gene analysis in each BC sample identified the top modulated genes and their frequency (**Figure 4b**). Two genes emerged as significantly modulated in most BC samples: *EMILIN1* and *COL3A1*. Both are related to TGF-β signaling, but in a different fashion. *COL3A1* is produced by fibroblasts in response to TGF-β activation^30^, whereas *EMILIN1* is an inhibitor of TGF-β signaling^31^. We were particularly interested in *EMILIN1* because its expression may modulate TGF-β activity and thus explain the selective CD8^+^ T-cell infiltration. A detailed comparison of *EMILIN1* expression in CD8^+^ T cell-low *versus* -high areas for each patient is provided in the **Figure S14**. Targeted analysis of the single-cell RNA-seq dataset reported by Wu et al.^23^ revealed that *EMILIN1* was a *bona fide* CAF gene, and was expressed only by myCAFs (**Figure 4c**). A more detailed analysis using the dataset reported by Kieffer et al.^25^ showed that *EMILIN1* was expressed by most CAF subpopulations, except IL-iCAFs. The strongest expression was observed in IFNγ-iCAFs, followed by ECM-myCAFs, IFNαβ-myCAFs and TGFβ-myCAFs (**Figure 4d**). Interestingly, wound-myCAFs, which showed the strongest expression of TGF-β signature genes (**Figure 4e**), displayed low *EMILIN1* expression.

**Figure 3:**
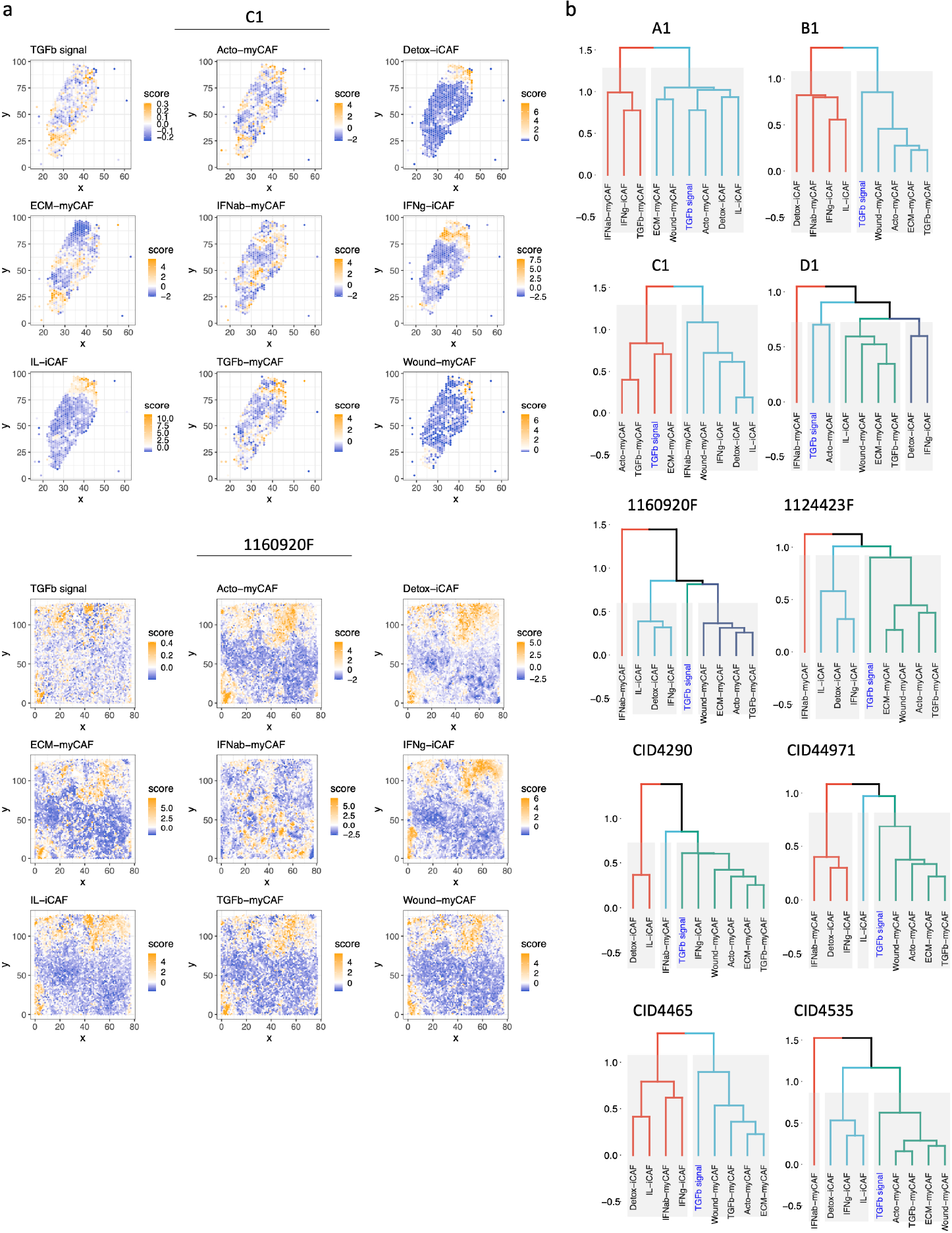
Spatial relationship of TGF-β signaling and CAF subpopulations in breast cancer. (**a**) Spatial distribution of genes implicated in TGF-β signaling (top) and spatial distribution of different CAF subpopulations in two breast cancer samples. The dendrogram (bottom, right) shows the spatial co-occurrence between CAF subpopulations and TGF-β signaling. (**b**) Dendrograms showing the co-occurrence of different CAF populations and TGF-β signaling in the other eight breast cancer samples.

**Figure 4:**
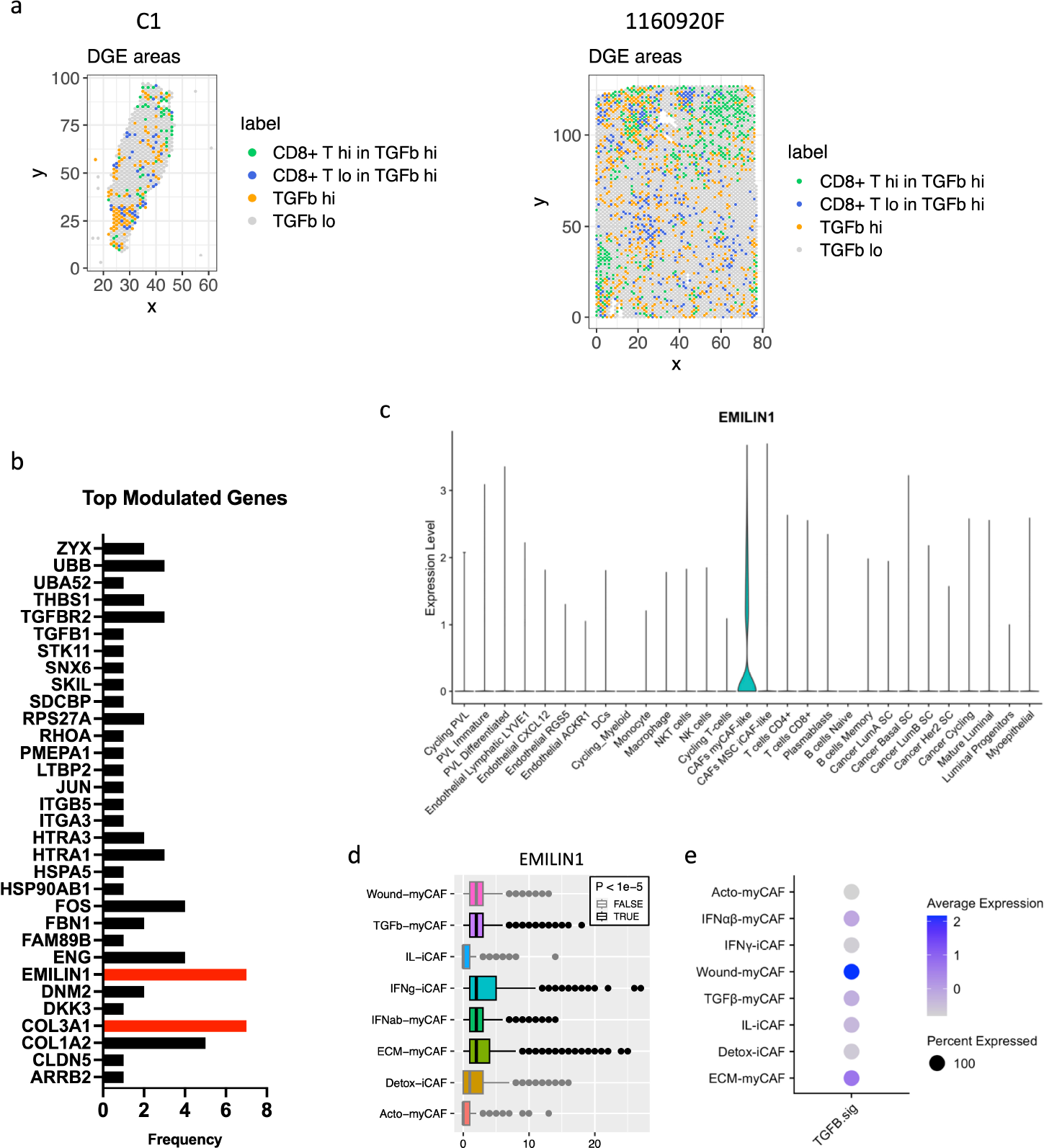
Differential gene expression analysis of areas with high in TGF-β signaling and with/without CD8^+^ T-cell exclusion. (**a**) Spatial distribution of areas with high *versus* low TGF-β signaling and presence/absence of CD8^+^ T cells in two breast cancer samples. (**b**) Differential gene expression analysis showing the frequency of top-modulated genes overexpressed in the areas where CD8^+^ T cells are present despite high TGF-β signaling. (**c**) *EMILIN1* expression in the indicated cell subpopulation (from the breast cancer atlas in **Figure 1a**). (**d**) *EMILIN1* expression in the indicated CAF subpopulations. (**e**) Upregulation of TGF-β signature genes in the indicated CAF subpopulations. The patient-wise statistical analysis of *EMILIN1* overexpression in CD8^+^ cells with high TGF-β signaling regions is provided in **Figure S14**

### Spatial modulation of EMILIN-1 expression coincides with CD8^+^ T-cell infiltration and is predictive of patient survival

To support the hypothesis that EMILIN1 expression is locally inhibiting TGF-β signaling, we monitored EMILIN-1 and TGFBI spatial expression by immunofluorescence analysis in 5 patients with BC. We selected TGFBI because this protein is a known TGF-β activity reporter in cancer^32^ and its expression is inversely correlated with CD8^+^ T-cell tumor infiltration^33^. TGFBI^high^ and EMILIN-1^high^ CAFs constituted two distinct cell populations (**Figure 5a**). CD8^+^ T cells were predominantly found in the EMILIN-rich areas, while they were excluded from regions with high TGFBI expression. As EMILIN-1 expression is limited to CAFs and EMILIN-1 functions as TGF-β activity suppressor^34^, we examined its expression by immunohistochemistry in 75 patients with BC and its relationship with CD8^+^ T-cell infiltration. We found that EMILIN-1 was clearly overexpressed in BC areas rich in infiltrating CD8^+^ T cells (**Figure 5b-c**). Moreover, EMILIN-1-rich areas had a significant proportion of Ki-67-negative cancer cells (**Figure 5c-e**), whereas many CD8^+^ T cells expressed Ki-67 (**Figure 5c**, yellow arrowheads). Lastly, survival analysis showed that high EMILIN-1 expression in BC was associated with increased survival (**Figure 5f**). The results of the multivariate analysis of EMILIN-1 expression in this patient cohort are in **Table S3**.

**Figure 5:**
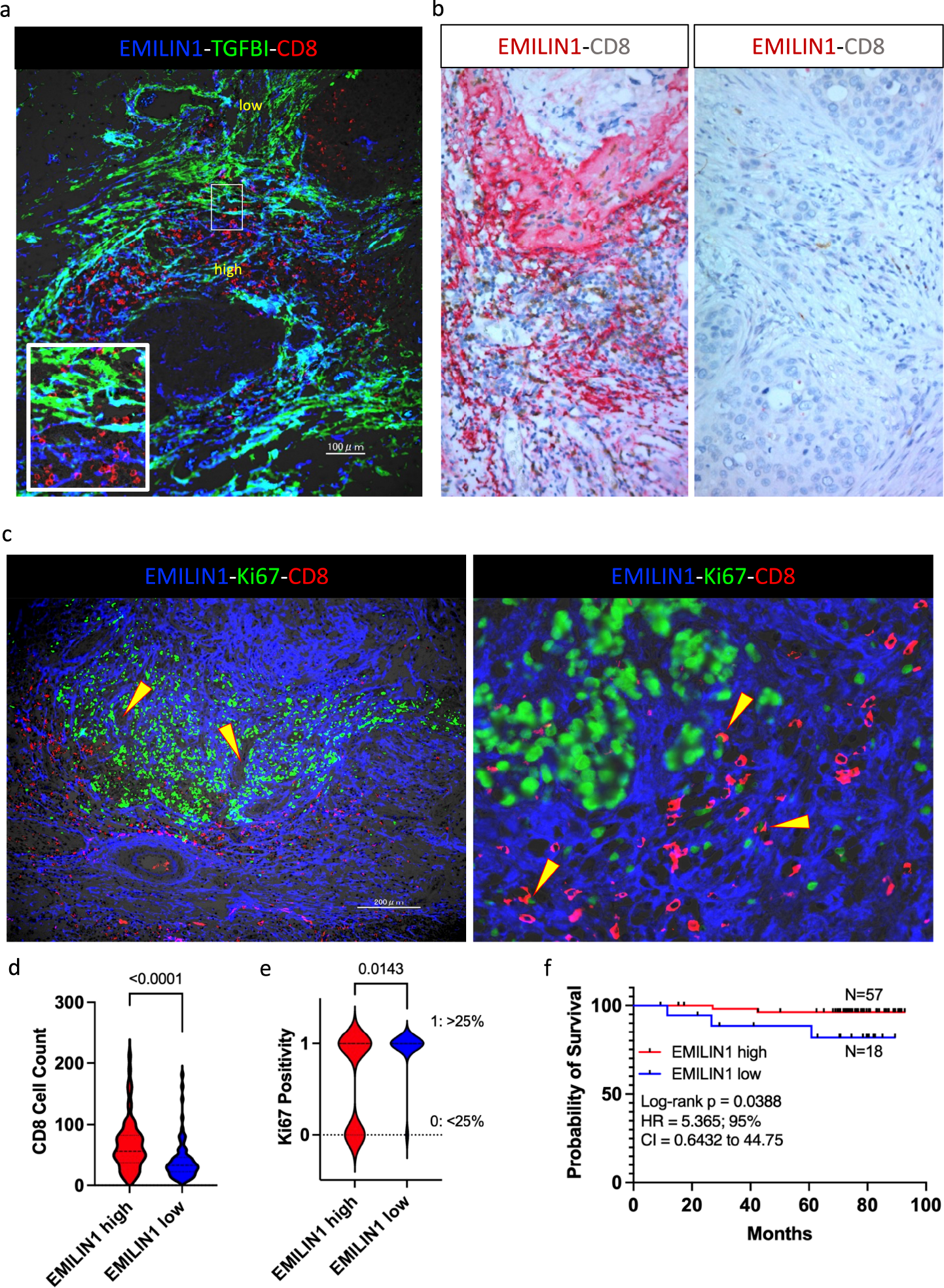
EMILIN-1 is a good prognostic marker in breast cancer. (a) Multiplexed immunofluorescence analysis displaying the localization of CD8^+^ T cells (red) and EMILIN-1 expression (blue) in CAFs in a representative breast cancer sample (N=5). Expression of TGFBI, a TGF-β signaling activity marker, was in green. (**b**) Multiplexed immunohistochemistry analysis showing examples of EMILIN-1 (red) and CD8 (brown) co-staining in breast cancer samples (N=75, all subtypes; for multivariate analysis see **Table S2**). (**c**) Low-and high-power views of the multiplexed immunofluorescence analysis displaying the localization of EMILIN-1, Ki-67 and CD8 in representative breast cancer samples (N=7). (**d-e**) Violin plots of CD8^+^ cell counts and Ki-67 positivity in areas of high *versus* low EMILIN-1 expression in breast cancer samples (N=75). (**f**) Survival analysis of patients with breast cancer (N=75) in function of EMILIN-1 expression level (high versus low) in the tumor. Maybe you should remind what you used as cut-off for high/low levels.

## DISCUSSION

ICIs have been a major breakthrough for the systemic treatment of some tumor types and patient subpopulations. The success of immunotherapy is influenced by the tumor immunological status and the infiltration of cytotoxic CD8^+^ T cells. Although the underlying mechanisms of their infiltration are poorly understood, CD8^+^ T cells are one of the most relevant effector cell types recruited by ICIs^35,36^. To shed additional light on CD8^+^ T-cell infiltration, the present study used comprehensive single-cell RNA-seq BC datasets to project the different cell populations spatially in BC tissue sections. In accordance with the literature, we found a clear, opposite, spatial correlation between proliferating cancer cells and CD8^+^ T cells in BC. It has also been shown that the tissue localization of CD8^+^ T cells is important for the patient outcome. CD8^+^ T cell presence in the tumor margin correlates with better clinical outcome^37,38^. However, we do not know which stromal parameters influence CD8^+^ T-cell composition, infiltration extent, activation, or exhaustion in BC. This limits our ability to turn immunologically cold tumors into hot tumors and in consequence subject them to effective ICI treatment. TGF-β signaling in the tumor stroma mediates this immunomodulatory process. TGF-β is a potent immune suppressor with direct effects on the proliferation, differentiation and survival of various immune cell sub-populations^39,40^. Experiments in mice suggest that TGF-β restricts CD8^+^ T-cell trafficking into tumors by suppressing CXCR3 expression^41^. These and other findings motivated the design of clinical trials to assess TGF-β blockade in combination with ICIs. However, the results were rather contrasted and surprisingly modest, in sharp contrast to the clear importance of TGF-β in tumor immunity^42^. The reason for this failure remains unclear, and might be related to the actual TGF-β activity, which is difficult to measure *in situ*. Indeed, TGF-β activity is modulated by factors secreted from the TME^29^. Therefore, TGF-β expression level may not be necessarily directly related to its activity level. To try to shed light into TGF-β activity, we spatially correlated the relationship between CD8^+^ T-cell tumor infiltration, individual cancer cell populations, and TGF-β activity in BC. CAFs are a TME cell type with specific features: they are major TGF-β producers in the tumor and among the largest modulators of its activity by expressing soluble matrix proteins that can efficiently inhibit this cytokine^43,44^. Therefore, it is not surprising that recent studies highlighted CAFs and some CAF subpopulations as key modulators of T-cell exclusion^45^. In the present study, spatial differences were mainly observed between the broad myCAF and iCAF subtypes, and myCAF were frequently spatially associated with higher TGF-β signaling. This is in line with previous studies in pancreatic cancer where such spatial heterogeneity was first reported^45,46^. A finer spatial distinction between all CAF subpopulations was not accessible, possibly due to limitations of the current spatial and single-cell transcriptomic data depth. Therefore, we performed differential analysis not based on individual CAF subpopulations but on regions with high TGF-β signaling and different degrees of CD8^+^ T-cell infiltration. This allowed us to propose a molecular explanation of the overlaps between TGF-β-driven myCAFs and areas of CD8^+^ T-cell infiltration, despite the known immunosuppressing effect of TGF-β. This analysis also highlighted a novel immune modulator protein, EMILIN-1, that was previously reported as a TGF-β inhibitor. We found that EMILIN-1 promotes CD8^+^ T-cell infiltration and is associated with better outcome in patients with BC. The results highlight the fact that different CAF populations cannot be simply categorized as immune-promoting or -suppressing cells and that their status is finely tuned by the expression of modulator genes. Such modulators can mitigate the activity of key cytokines, such as TGF-β. Moreover, this finding suggests that CAF subpopulations can be largely regarded as cell programing states, probably with few exceptions. Such exceptions may occur in organs/tissues where different CAF sources are possible because of the intrinsic presence of different fibroblast-like cells (e.g. stellate cells in liver). Viewing CAF heterogeneity as cell states rather than actual subpopulations implies that harnessing CAFs for therapy would require their re-programing rather than the elimination of a specific CAF subpopulation. In this regard the current study highlights EMILIN-1 as an important determinant of CAF anti-tumor program. Future studies should elucidate how EMILIN-1 expression is modulated and how it could be upregulated in CAFs.

## ACKNOWLEDGEMENTS

The authors want to thank Ms. Mariko Nakamura, Kao Abe, Kei Masuda, Kumiko Sudo, Yukiko Suto, Kyoko Miyoshi, Miyoko Suzuki, Chiho Noguchi, Saori Suto, Sayaka Okada, and Harumi Kanai for experimental assistance. This work was supported by grants from Gunma University. A.T. is supported by a LabEx MabImprove Starting Grant. J.C. was supported by a Fondation ARC grant PJA 20141201975. No funding body had any role in the study design, data collection and analysis, decision to publish, or preparation of the manuscript.

## Notes

### Competing Interest Statement

The authors have declared no competing interest.

