## Supplemental Figures for "Cancer-associated Fibroblast Spatial Heterogeneity and *EMILIN1* Expression in Cancer Stroma Modulate TGF-β Activity and CD8^+^ T-Cell Infiltration in Breast Cancer"

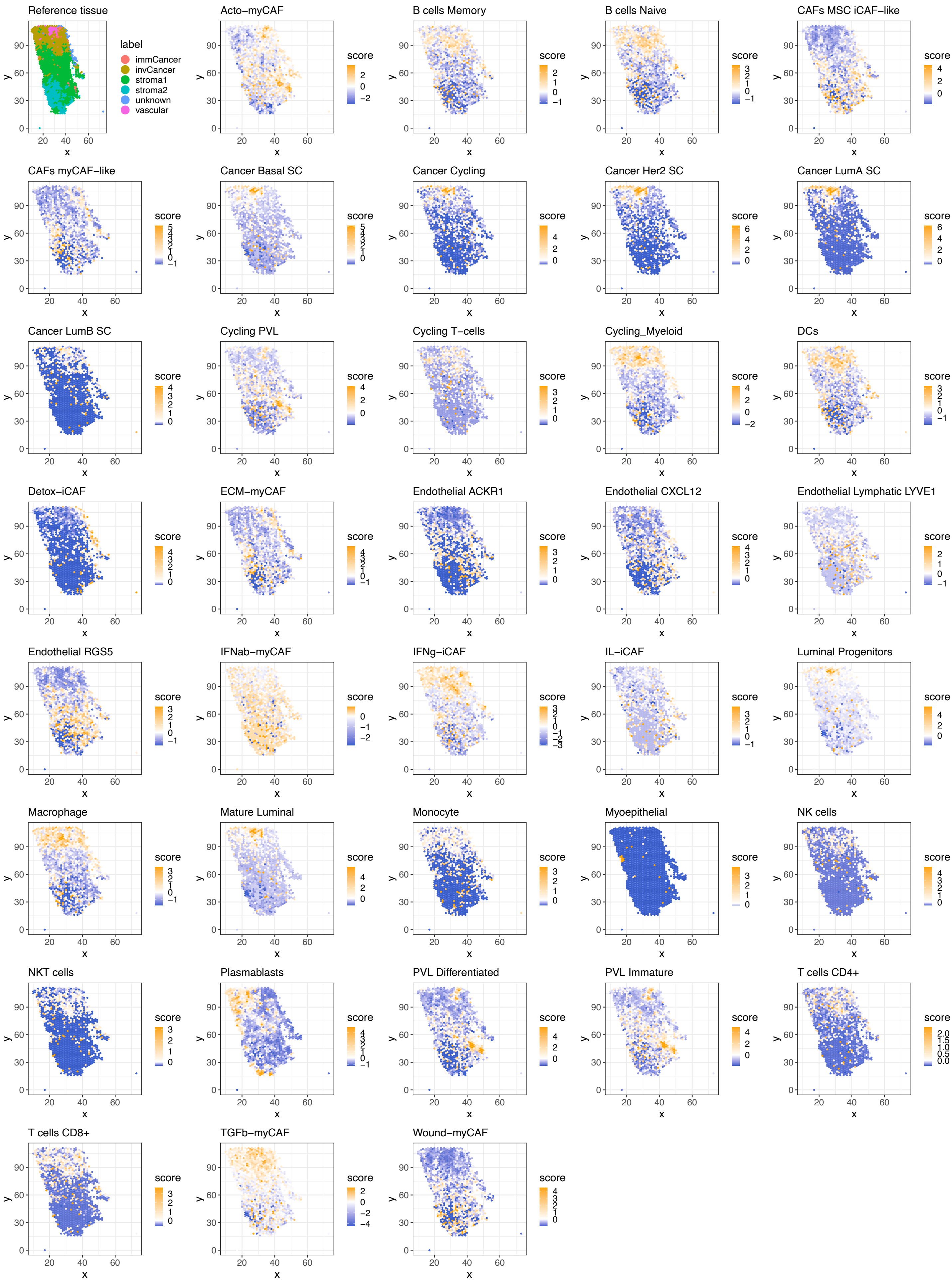

Reference tissue
label
immCancer
invCancer
stroma1
stroma2
unknown
vascular
Acto−myCAF
B cells Memory
B cells Naive
CAFs MSC iCAF−like
CAFs myCAF−like
Cancer Basal SC
Cancer Cycling
Cancer Her2 SC
Cancer LumA SC
Cancer LumB SC
Cycling PVL
Cycling T−cells
Cycling_Myeloid
DCs
Detox−iCAF
ECM−myCAF
Endothelial ACKR1
Endothelial CXCL12
Endothelial Lymphatic LYVE1
Endothelial RGS5
IFNab−myCAF
IFNg−iCAF
IL−iCAF
Luminal Progenitors
Macrophage
Mature Luminal
Monocyte
Myoepithelial
NK cells
NKT cells
Plasmablasts
PVL Differentiated
PVL Immature
T cells CD4+
T cells CD8+
TGFb−myCAF
Wound−myCAF
score
x
y


Figure S1

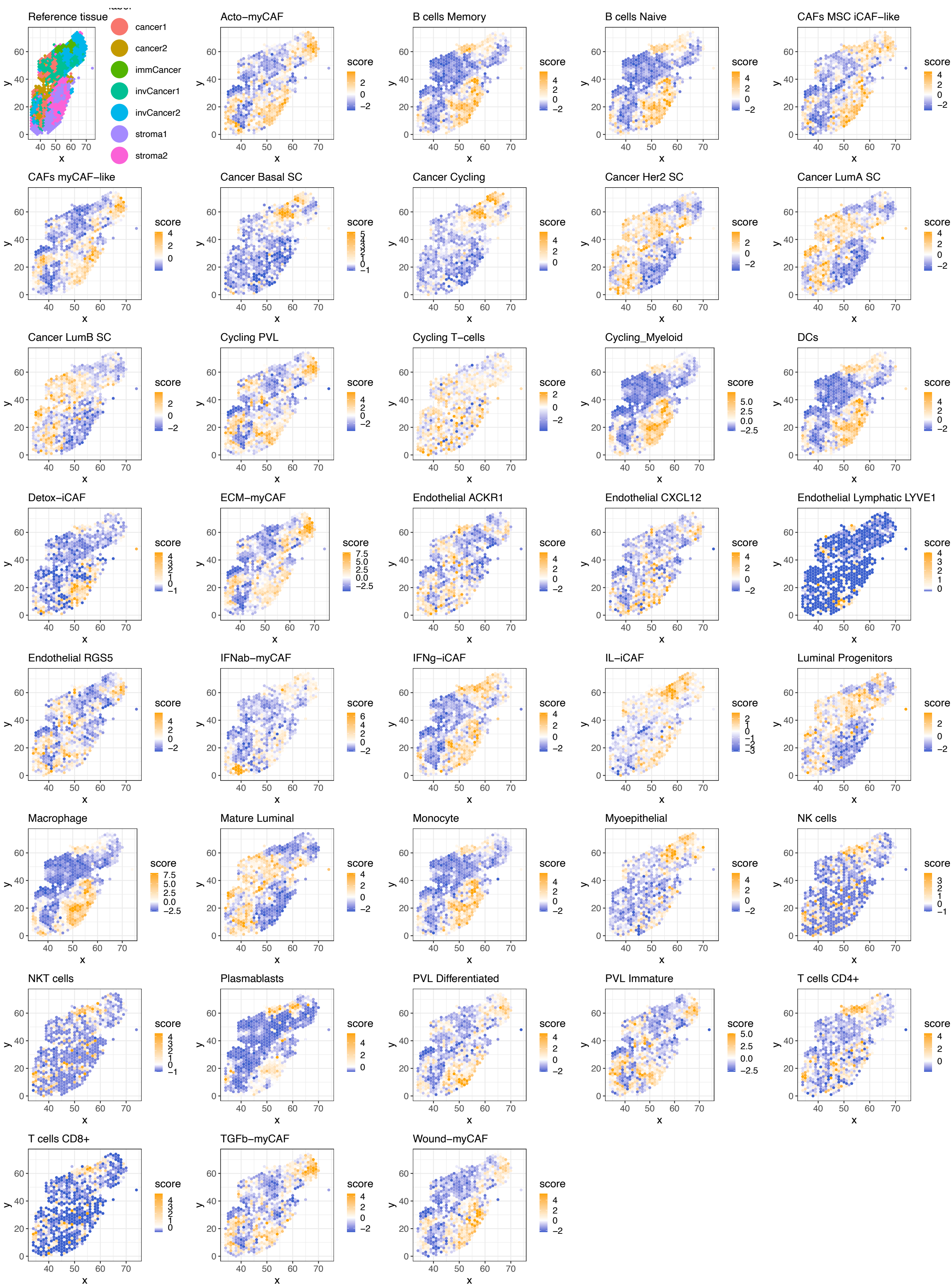
Reference tissue
label
cancer1
cancer2
immCancer
invCancer1
invCancer2
stroma1
stroma2
Acto−myCAF
B cells Memory
B cells Naive
CAFs MSC iCAF−like
CAFs myCAF−like
Cancer Basal SC
Cancer Cycling
Cancer Her2 SC
Cancer LumA SC
Cancer LumB SC
Cycling PVL
Cycling T−cells
Cycling_Myeloid
DCs
Detox−iCAF
ECM−myCAF
Endothelial ACKR1
Endothelial CXCL12
Endothelial Lymphatic LYVE1
Endothelial RGS5
IFNab−myCAF
IFNg−iCAF
IL−iCAF
Luminal Progenitors
Macrophage
Mature Luminal
Monocyte
Myoepithelial
NK cells
NKT cells
Plasmablasts
PVL Differentiated
PVL Immature
T cells CD4+
T cells CD8+
TGFb−myCAF
Wound−myCAF
score
x
y

Figure S2

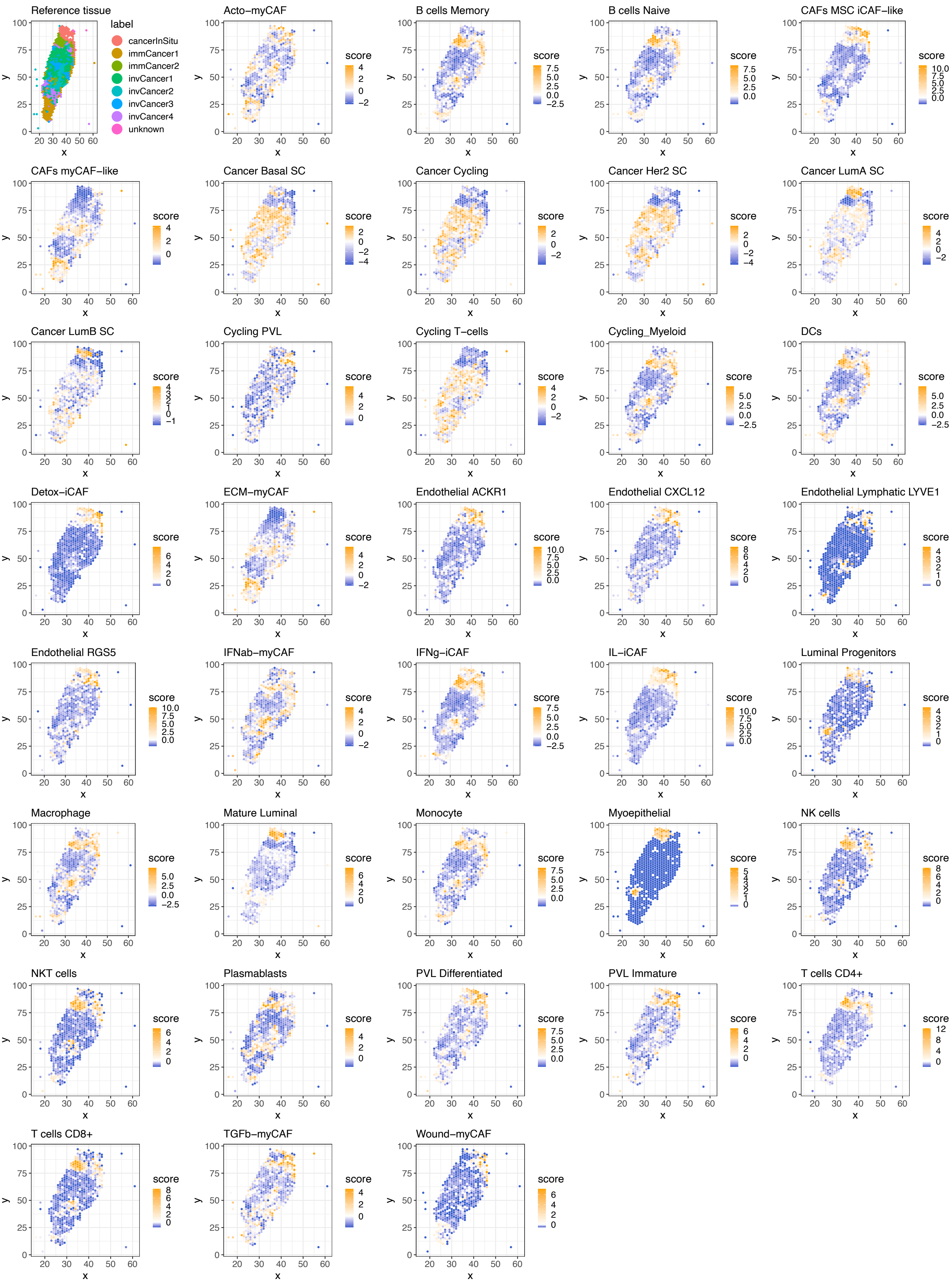
Reference tissue
label
cancerInSitu
immCancer1
immCancer2
invCancer1
invCancer2
invCancer3
invCancer4
unknown
Acto−myCAF
B cells Memory
B cells Naive
CAFs MSC iCAF−like
CAFs myCAF−like
Cancer Basal SC
Cancer Cycling
Cancer Her2 SC
Cancer LumA SC
Cancer LumB SC
Cycling PVL
Cycling T−cells
Cycling_Myeloid
DCs
Detox−iCAF
ECM−myCAF
Endothelial ACKR1
Endothelial CXCL12
Endothelial Lymphatic LYVE1
Endothelial RGS5
IFNab−myCAF
IFNg−iCAF
IL−iCAF
Luminal Progenitors
Macrophage
Mature Luminal
Monocyte
Myoepithelial
NK cells
NKT cells
Plasmablasts
PVL Differentiated
PVL Immature
T cells CD4+
T cells CD8+
TGFb−myCAF
Wound−myCAF
score
x
y

Figure S3

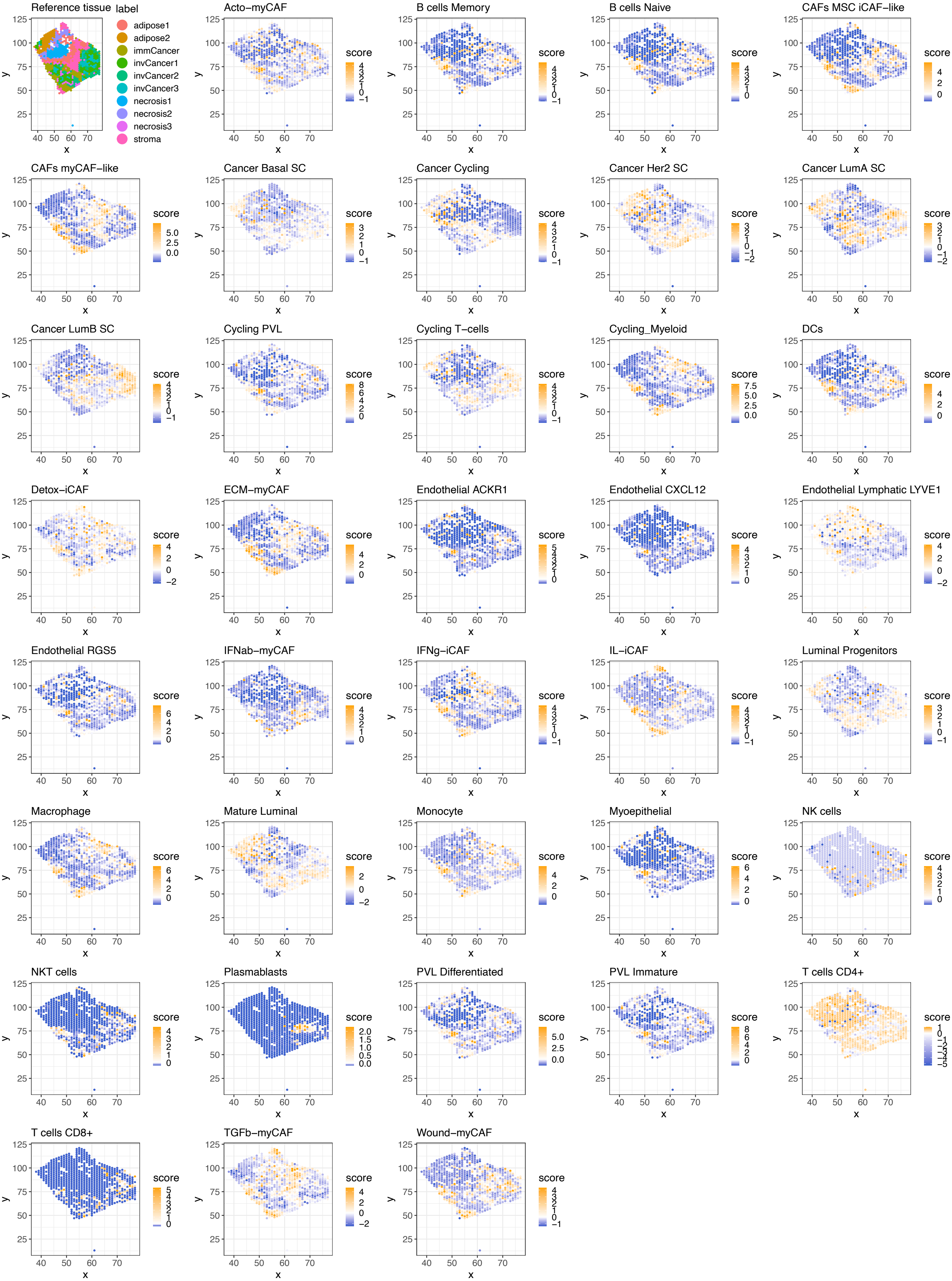
Reference tissue
label
adipose1
adipose2
immCancer
invCancer1
invCancer2
invCancer3
necrosis1
necrosis2
necrosis3
stroma
Acto−myCAF
B cells Memory
B cells Naive
CAFs MSC iCAF−like
CAFs myCAF−like
Cancer Basal SC
Cancer Cycling
Cancer Her2 SC
Cancer LumA SC
Cancer LumB SC
Cycling PVL
Cycling T−cells
Cycling_Myeloid
DCs
Detox−iCAF
ECM−myCAF
Endothelial ACKR1
Endothelial CXCL12
Endothelial Lymphatic LYVE1
Endothelial RGS5
IFNab−myCAF
IFNg−iCAF
IL−iCAF
Luminal Progenitors
Macrophage
Mature Luminal
Monocyte
Myoepithelial
NK cells
NKT cells
Plasmablasts
PVL Differentiated
PVL Immature
T cells CD4+
T cells CD8+
TGFb−myCAF
Wound−myCAF
score
x
y

Figure S4

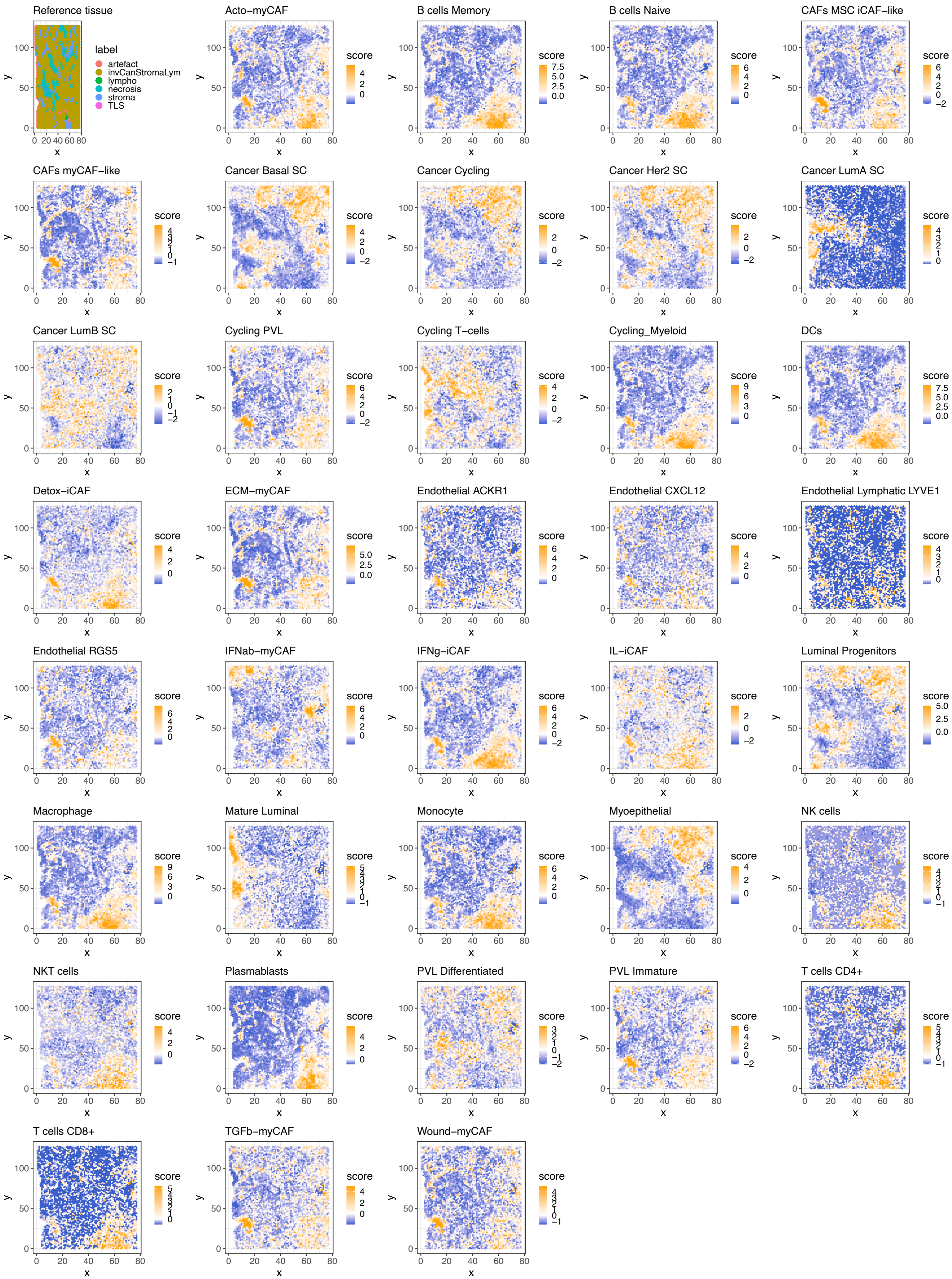
Reference tissue
label
artefact
invCanStromaLym
lympho
necrosis
stroma
TLS
Acto−myCAF
B cells Memory
B cells Naive
CAFs MSC iCAF−like
CAFs myCAF−like
Cancer Basal SC
Cancer Cycling
Cancer Her2 SC
Cancer LumA SC
Cancer LumB SC
Cycling PVL
Cycling T−cells
Cycling_Myeloid
DCs
Detox−iCAF
ECM−myCAF
Endothelial ACKR1
Endothelial CXCL12
Endothelial Lymphatic LYVE1
Endothelial RGS5
IFNab−myCAF
IFNg−iCAF
IL−iCAF
Luminal Progenitors
Macrophage
Mature Luminal
Monocyte
Myoepithelial
NK cells
NKT cells
Plasmablasts
PVL Differentiated
PVL Immature
T cells CD4+
T cells CD8+
TGFb−myCAF
Wound−myCAF
score
x
y

Figure S5

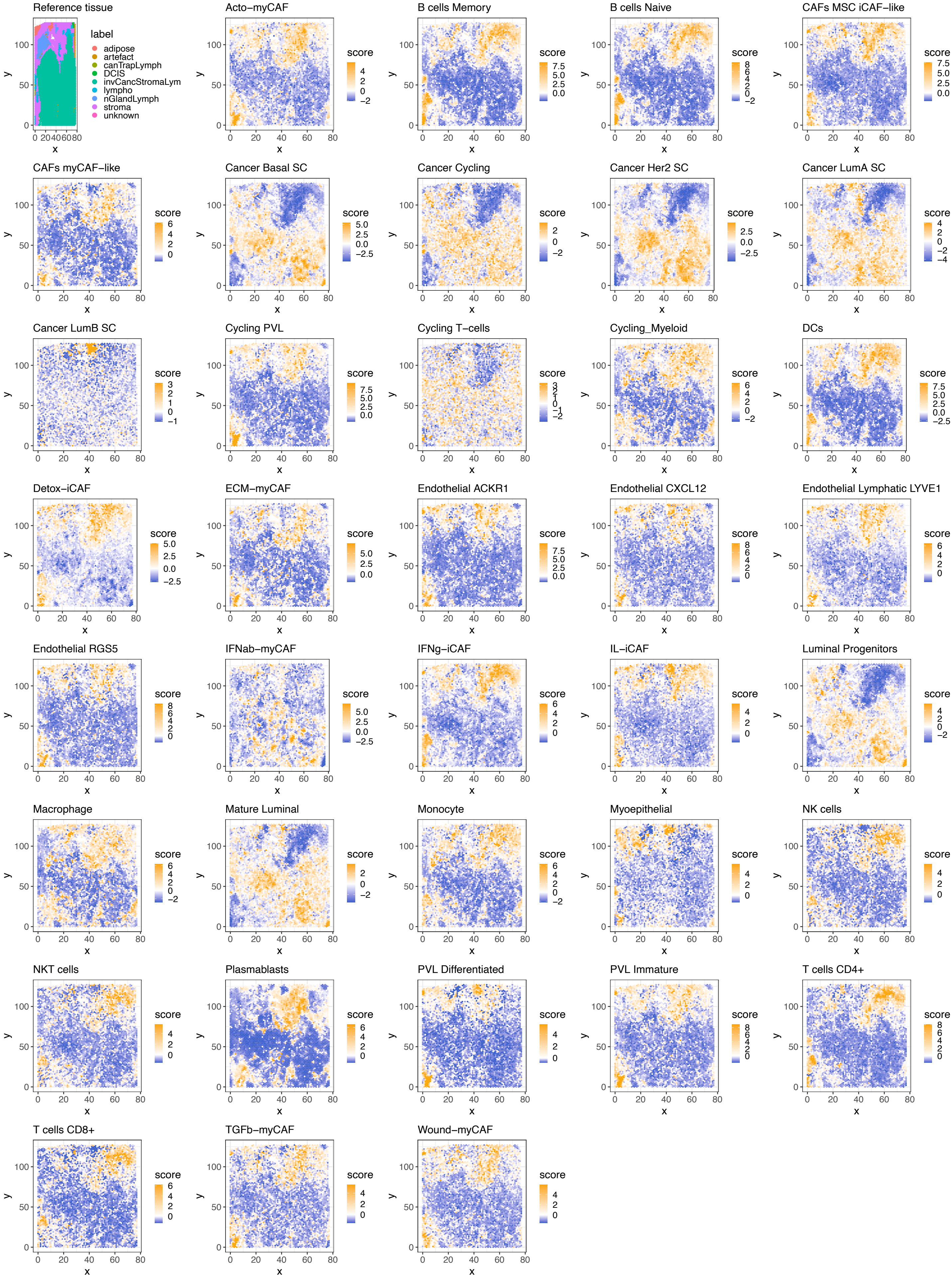
Reference tissue
label
adipose
artefact
canTrapLymph
DCIS
invCancStromaLym
lympho
nGlandLymph
stroma
unknown
Acto−myCAF
B cells Memory
B cells Naive
CAFs MSC iCAF−like
CAFs myCAF−like
Cancer Basal SC
Cancer Cycling
Cancer Her2 SC
Cancer LumA SC
Cancer LumB SC
Cycling PVL
Cycling T−cells
Cycling_Myeloid
DCs
Detox−iCAF
ECM−myCAF
Endothelial ACKR1
Endothelial CXCL12
Endothelial Lymphatic LYVE1
Endothelial RGS5
IFNab−myCAF
IFNg−iCAF
IL−iCAF
Luminal Progenitors
Macrophage
Mature Luminal
Monocyte
Myoepithelial
NK cells
NKT cells
Plasmablasts
PVL Differentiated
PVL Immature
T cells CD4+
T cells CD8+
TGFb−myCAF
Wound−myCAF
score
x
y

Figure S6

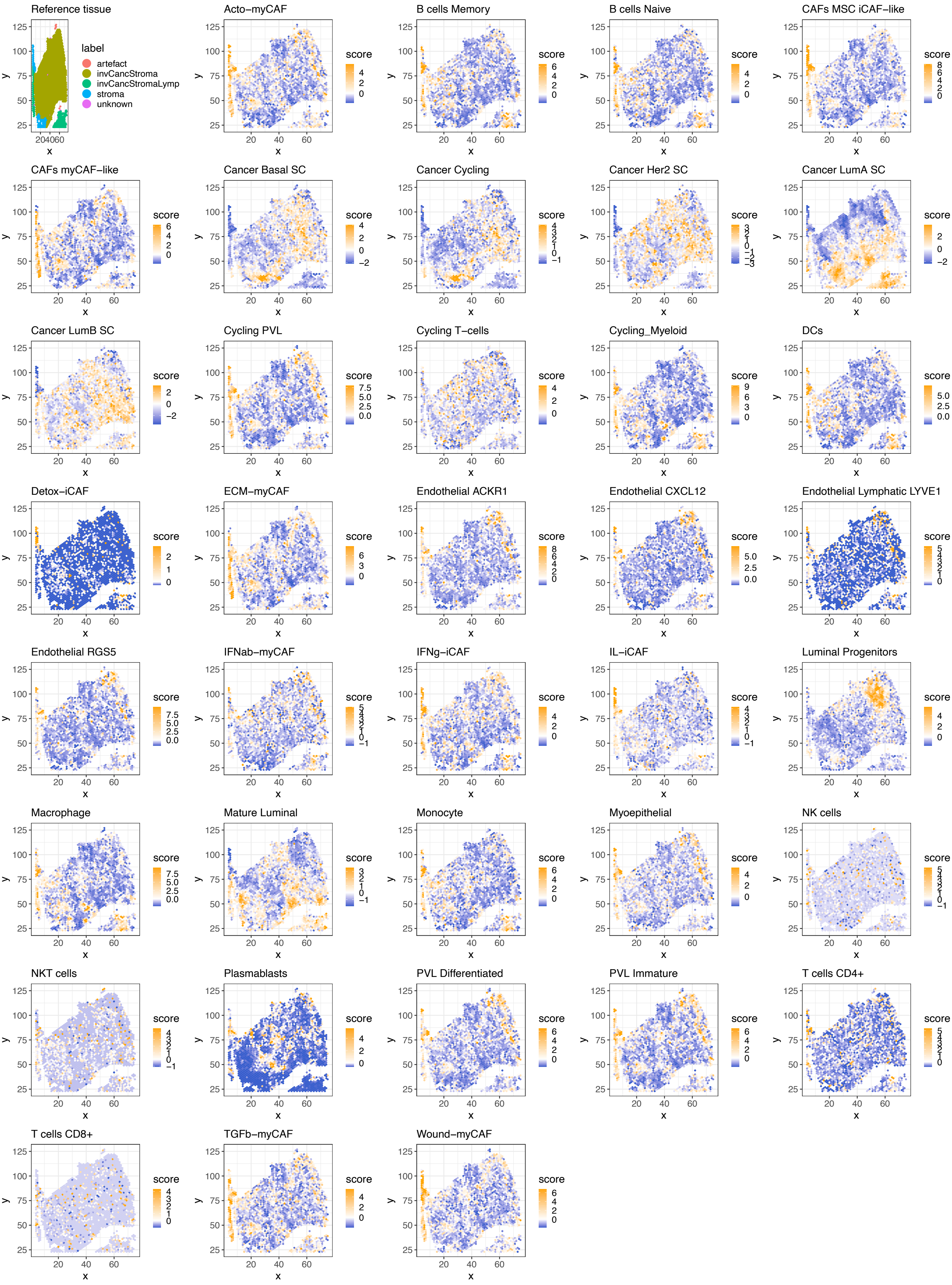
Reference tissue
label
artefact
invCancStroma
invCancStromaLymp
stroma
unknown
Acto−myCAF
B cells Memory
B cells Naive
CAFs MSC iCAF−like
CAFs myCAF−like
Cancer Basal SC
Cancer Cycling
Cancer Her2 SC
Cancer LumA SC
Cancer LumB SC
Cycling PVL
Cycling T−cells
Cycling_Myeloid
DCs
Detox−iCAF
ECM−myCAF
Endothelial ACKR1
Endothelial CXCL12
Endothelial Lymphatic LYVE1
Endothelial RGS5
IFNab−myCAF
IFNg−iCAF
IL−iCAF
Luminal Progenitors
Macrophage
Mature Luminal
Monocyte
Myoepithelial
NK cells
NKT cells
Plasmablasts
PVL Differentiated
PVL Immature
T cells CD4+
T cells CD8+
TGFb−myCAF
Wound−myCAF
score
x
y

Figure S7

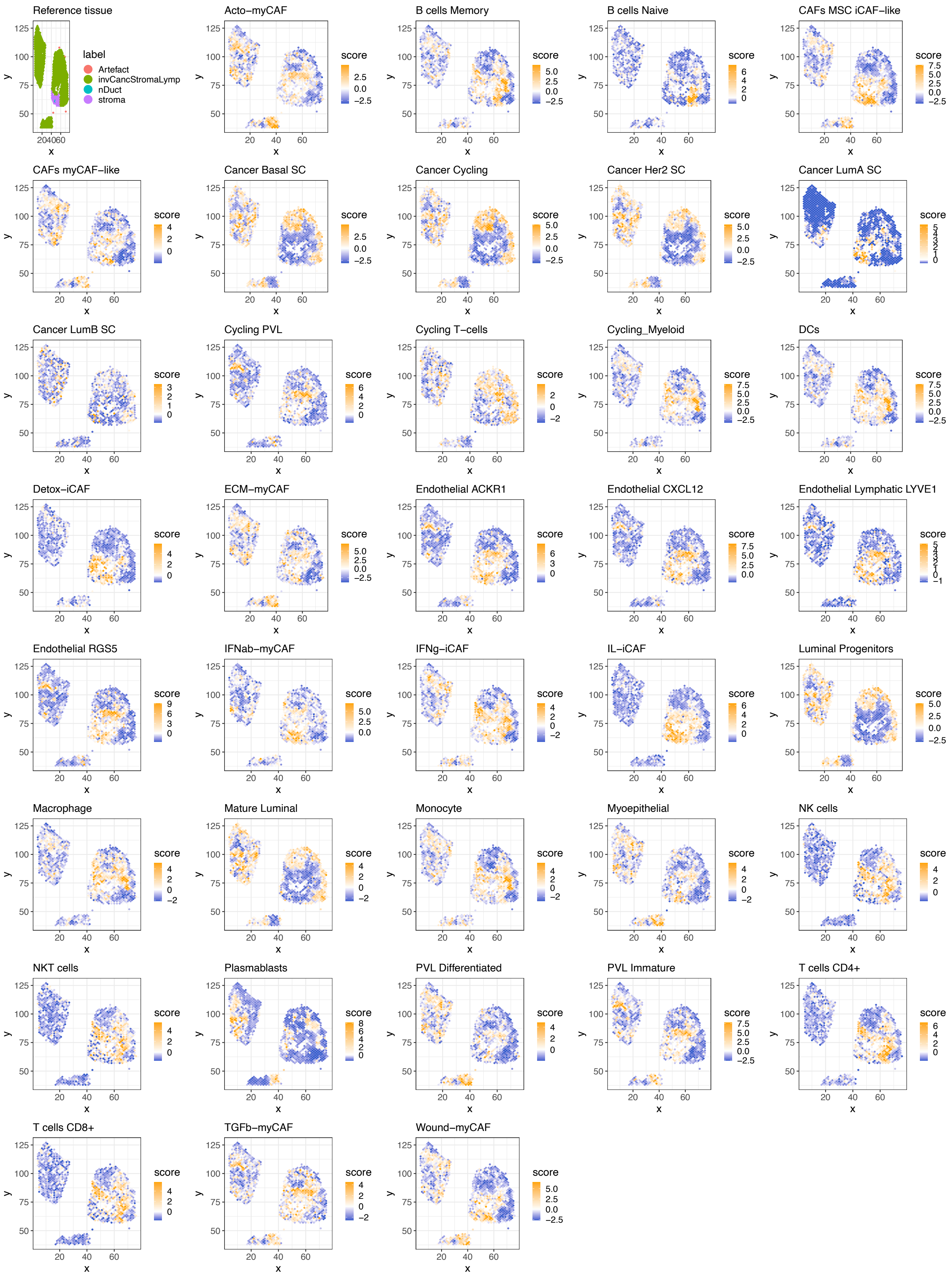
Reference tissue
label
Artefact
invCancStromaLymp
nDuct
stroma
Acto−myCAF
B cells Memory
B cells Naive
CAFs MSC iCAF−like
CAFs myCAF−like
Cancer Basal SC
Cancer Cycling
Cancer Her2 SC
Cancer LumA SC
Cancer LumB SC
Cycling PVL
Cycling T−cells
Cycling_Myeloid
DCs
Detox−iCAF
ECM−myCAF
Endothelial ACKR1
Endothelial CXCL12
Endothelial Lymphatic LYVE1
Endothelial RGS5
IFNab−myCAF
IFNg−iCAF
IL−iCAF
Luminal Progenitors
Macrophage
Mature Luminal
Monocyte
Myoepithelial
NK cells
NKT cells
Plasmablasts
PVL Differentiated
PVL Immature
T cells CD4+
T cells CD8+
TGFb−myCAF
Wound−myCAF
score
x
y

Figure S8

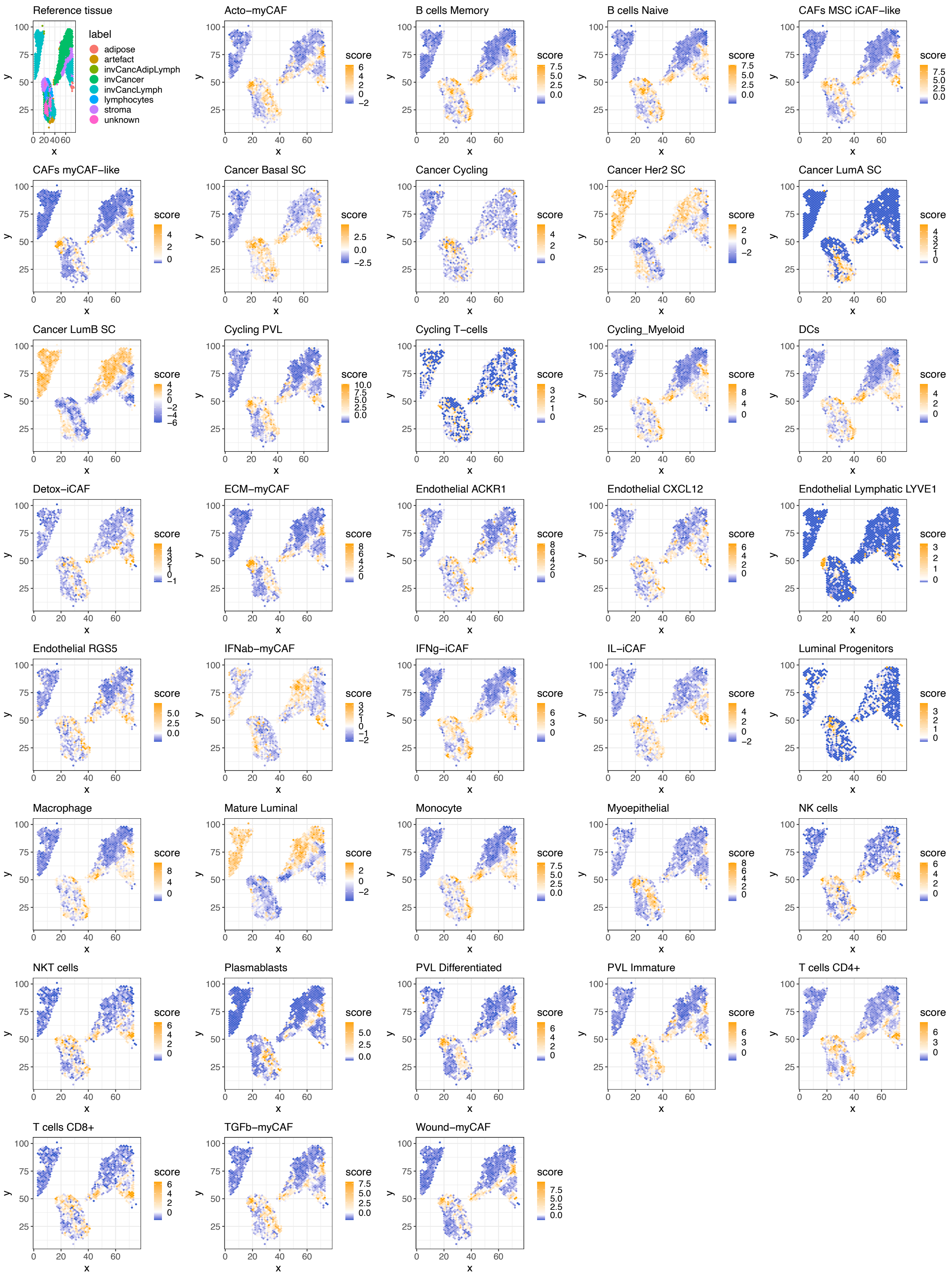
Reference tissue
label
adipose
artefact
invCancAdipLymph
invCancer
invCancLymph
lymphocytes
stroma
unknown
Acto−myCAF
B cells Memory
B cells Naive
CAFs MSC iCAF−like
CAFs myCAF−like
Cancer Basal SC
Cancer Cycling
Cancer Her2 SC
Cancer LumA SC
Cancer LumB SC
Cycling PVL
Cycling T−cells
Cycling_Myeloid
DCs
Detox−iCAF
ECM−myCAF
Endothelial ACKR1
Endothelial CXCL12
Endothelial Lymphatic LYVE1
Endothelial RGS5
IFNab−myCAF
IFNg−iCAF
IL−iCAF
Luminal Progenitors
Macrophage
Mature Luminal
Monocyte
Myoepithelial
NK cells
NKT cells
Plasmablasts
PVL Differentiated
PVL Immature
T cells CD4+
T cells CD8+
TGFb−myCAF
Wound−myCAF
score
x
y

Figure S9

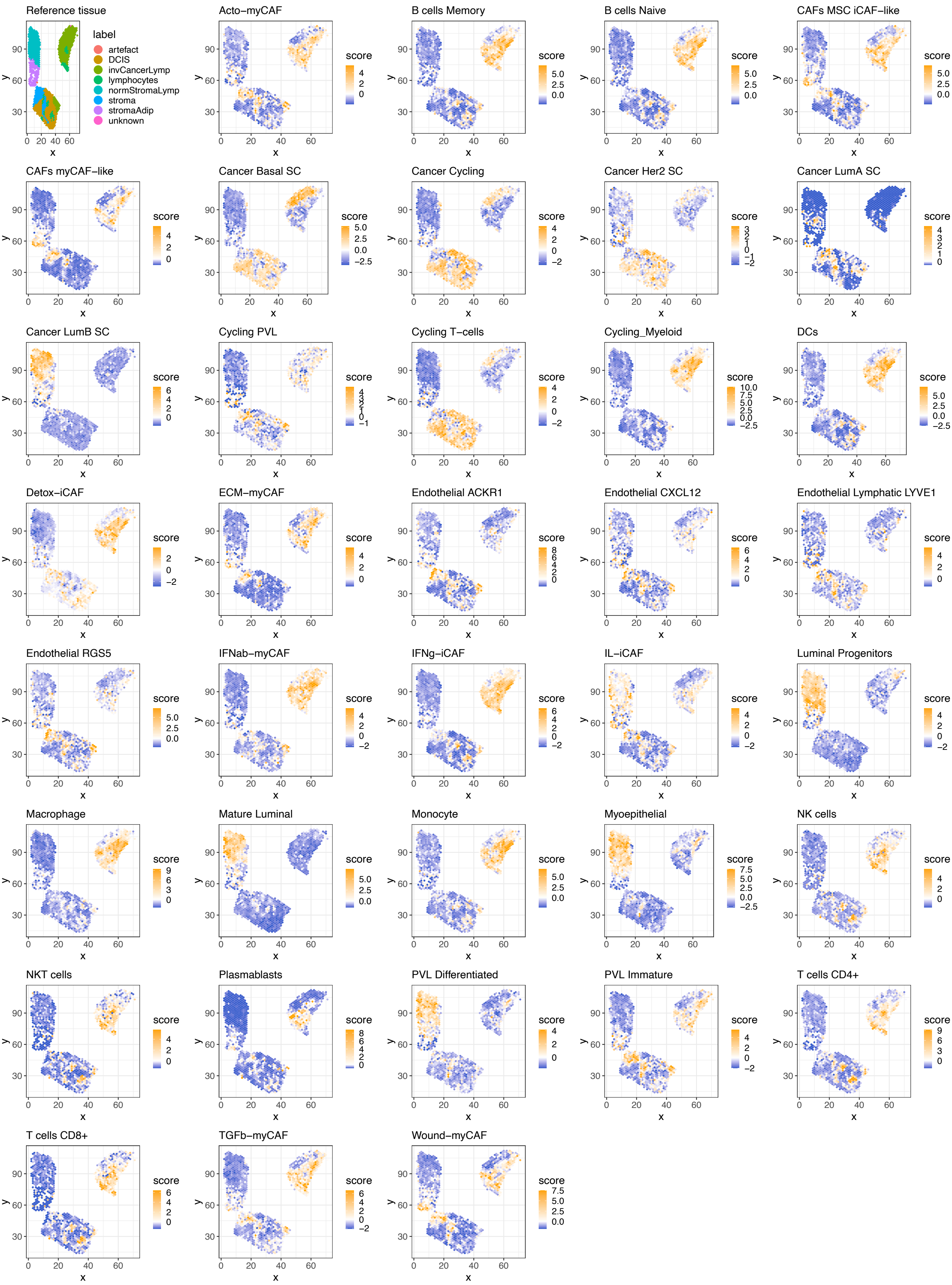
Reference tissue
label
artefact
DCIS
invCancerLymp
lymphocytes
normStromaLymp
stroma
stromaAdip
unknown
Acto−myCAF
B cells Memory
B cells Naive
CAFs MSC iCAF−like
CAFs myCAF−like
Cancer Basal SC
Cancer Cycling
Cancer Her2 SC
Cancer LumA SC
Cancer LumB SC
Cycling PVL
Cycling T−cells
Cycling_Myeloid
DCs
Detox−iCAF
ECM−myCAF
Endothelial ACKR1
Endothelial CXCL12
Endothelial Lymphatic LYVE1
Endothelial RGS5
IFNab−myCAF
IFNg−iCAF
IL−iCAF
Luminal Progenitors
Macrophage
Mature Luminal
Monocyte
Myoepithelial
NK cells
NKT cells
Plasmablasts
PVL Differentiated
PVL Immature
T cells CD4+
T cells CD8+
TGFb−myCAF
Wound−myCAF
score
x
y

Figure S10

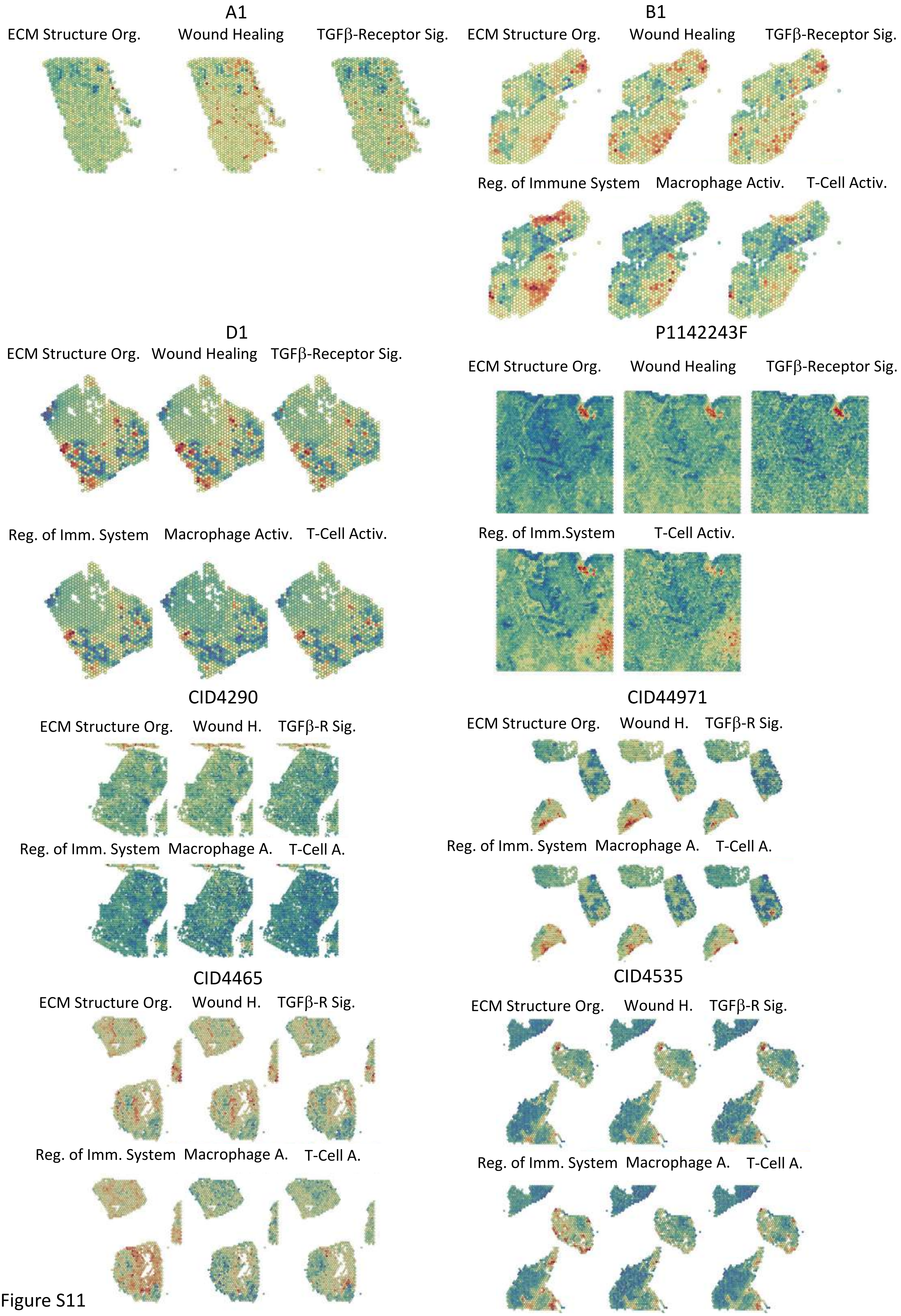

A1
ECM Structure Org.
Wound Healing
TGFβ-Receptor Sig.
B1
ECM Structure Org.
Wound Healing
TGFβ-Receptor Sig.
Reg. of Immune System
Macrophage Activ.
T-Cell Activ.
D1
ECM Structure Org.
Wound Healing
TGFβ-Receptor Sig.
Reg. of Imm. System
Macrophage Activ.
T-Cell Activ.
P1142243F
ECM Structure Org.
Wound Healing
TGFβ-Receptor Sig.
Reg. of Imm.System
T-Cell Activ.
CID4290
ECM Structure Org.
Wound H.
TGFβ-R Sig.
Reg. of Imm. System
Macrophage A.
T-Cell A.
CID44971
ECM Structure Org.
Wound H.
TGFβ-R Sig.
Reg. of Imm. System
Macrophage A.
T-Cell A.
CID4465
ECM Structure Org.
Wound H.
TGFβ-R Sig.
Reg. of Imm. System
Macrophage A.
T-Cell A.
CID4535
ECM Structure Org.
Wound H.
TGFβ-R Sig.
Reg. of Imm. System
Macrophage A.
T-Cell A.


Figure S11

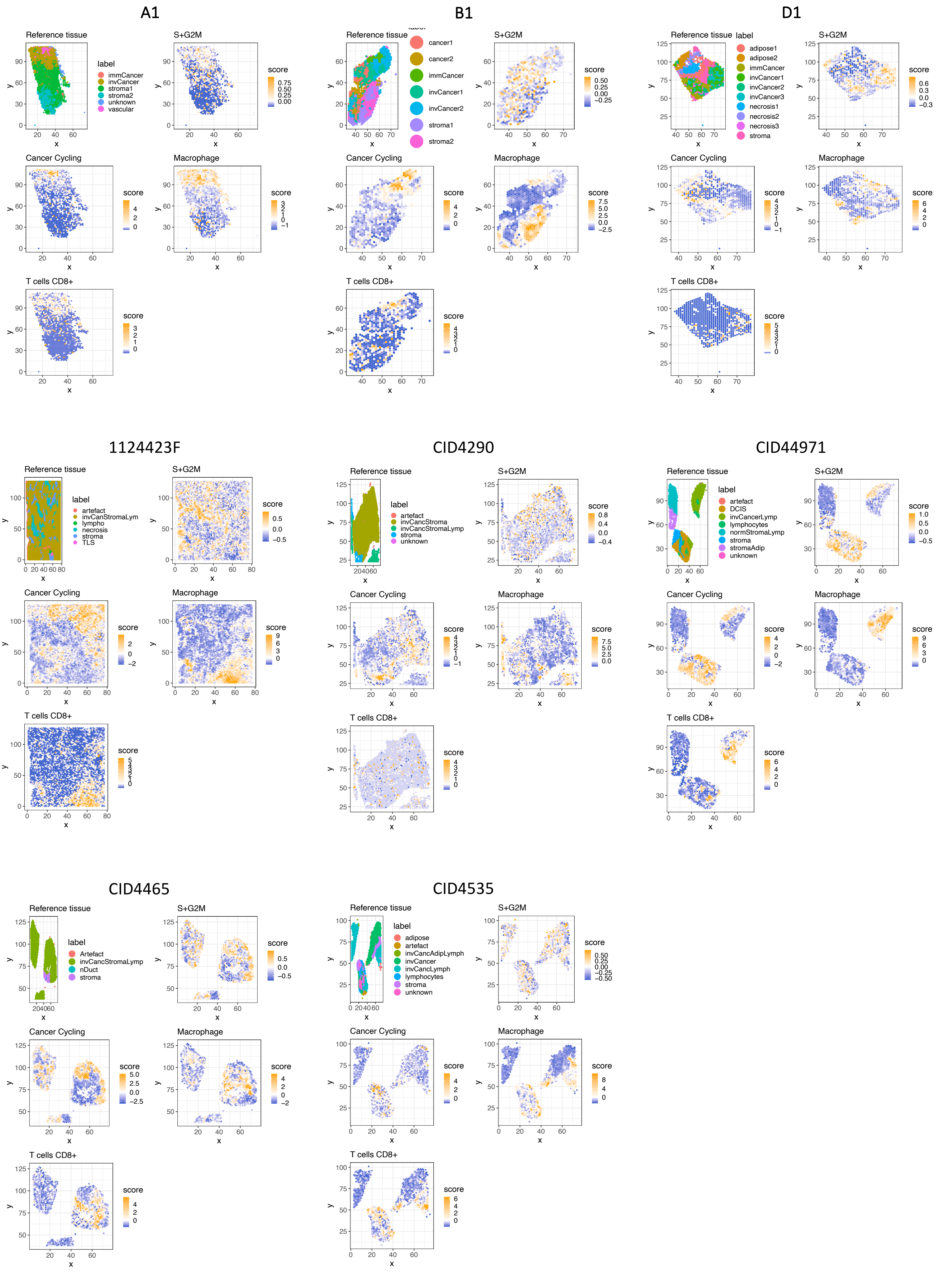
A1
Reference tissue
label
immCancer
invCancer
stroma1
stroma2
unknown
vascular
S+G2M
Cancer Cycling
Macrophage
T cells CD8+
score
B1
Reference tissue
label
cancer1
cancer2
immCancer
invCancer1
invCancer2
stroma1
stroma2
S+G2M
Cancer Cycling
Macrophage
T cells CD8+
D1
Reference tissue
label
adipose1
adipose2
immCancer
invCancer1
invCancer2
invCancer3
necrosis1
necrosis2
necrosis3
stroma
S+G2M
Cancer Cycling
Macrophage
T cells CD8+
1124423F
Reference tissue
label
artefact
invCanStromaLym
lympho
necrosis
stroma
TLS
S+G2M
Cancer Cycling
Macrophage
T cells CD8+
CID4290
Reference tissue
label
artefact
invCancStroma
invCancStromaLymp
stroma
unknown
S+G2M
Cancer Cycling
Macrophage
T cells CD8+
CID44971
Reference tissue
label
artefact
DCIS
invCancerLymp
lymphocytes
normStromaLymp
stroma
stromaAdip
unknown
S+G2M
Cancer Cycling
Macrophage
T cells CD8+
CID4465
Reference tissue
label
Artefact
invCancStromaLymp
nDuct
stroma
S+G2M
Cancer Cycling
Macrophage
T cells CD8+
CID4535
Reference tissue
label
adipose
artefact
invCancAdipLymph
invCancer
invCancLymph
lymphocytes
stroma
unknown
S+G2M
Cancer Cycling
Macrophage
T cells CD8+
x
y
score

Figure S12

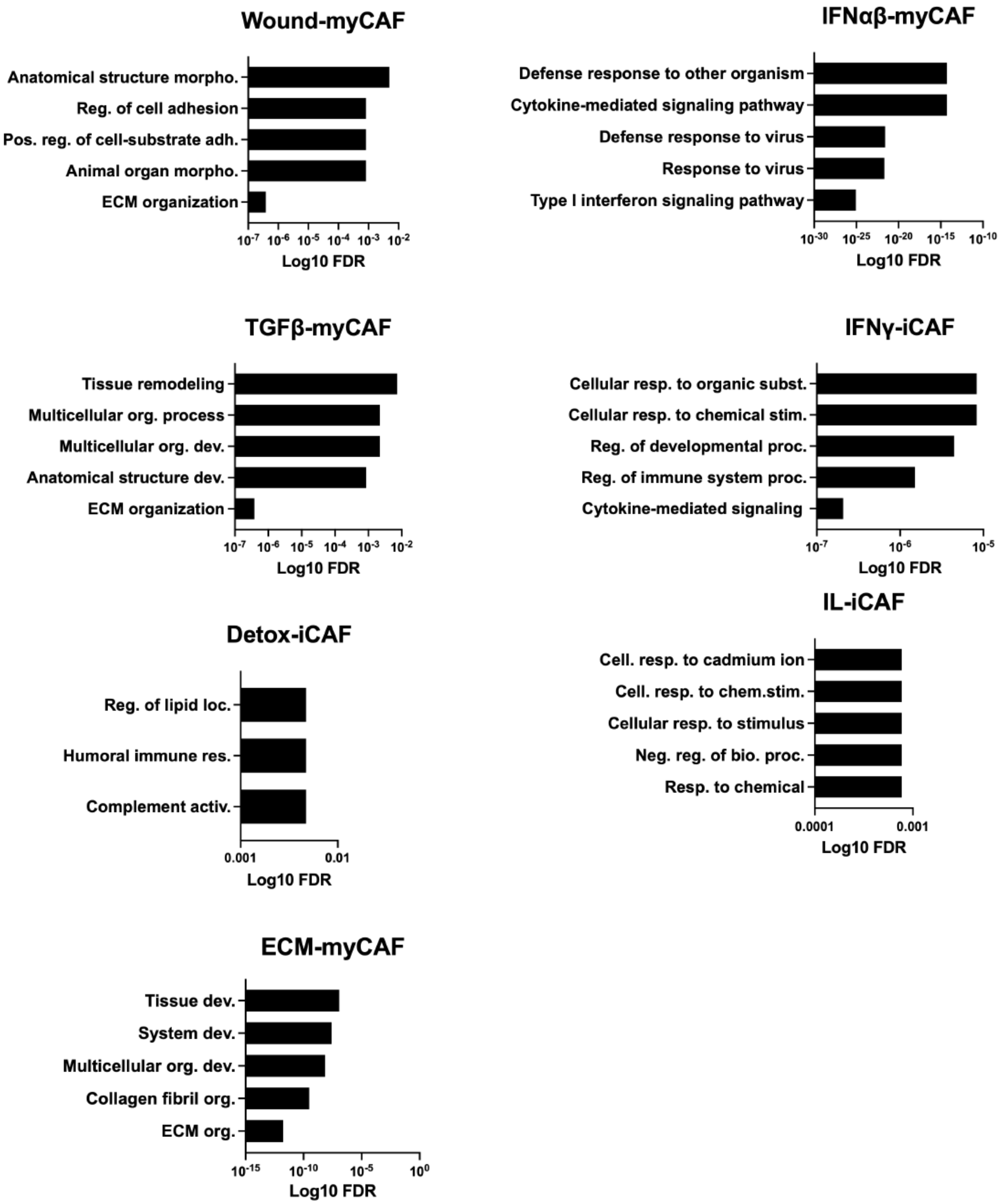

Wound-myCAF
Anatomical structure morpho.
Reg. of cell adhesion
Pos. reg. of cell-substrate adh.
Animal organ morpho.
ECM organization
10-7 10-6 10-5 10-4 10-3 10-2
Log10 FDR
IFNαβ-myCAF
Defense response to other organism
Cytokine-mediated signaling pathway
Defense response to virus
Response to virus
Type I interferon signaling pathway
10-30 10-25 10-20 10-15 10-10
Log10 FDR
TGFβ-myCAF
Tissue remodeling
Multicellular org. process
Multicellular org. dev.
Anatomical structure dev.
ECM organization
10-7 10-6 10-5 10-4 10-3 10-2
Log10 FDR
IFNγ-iCAF
Cellular resp. to organic subst.
Cellular resp. to chemical stim.
Reg. of developmental proc.
Reg. of immune system proc.
Cytokine-mediated signaling
10-7 10-6 10-5
Log10 FDR
IL-iCAF
Cell. resp. to cadmium ion
Cell. resp. to chem.stim.
Cellular resp. to stimulus
Neg. reg. of bio. proc.
Resp. to chemical
0.0001 0.001
Log10 FDR
Detox-iCAF
Reg. of lipid loc.
Humoral immune res.
Complement activ.
0.001 0.01
Log10 FDR
ECM-myCAF
Tissue dev.
System dev.
Multicellular org. dev.
Collagen fibril org.
ECM org.
10-15 10-10 10-5 100
Log10 FDR


Figure S13

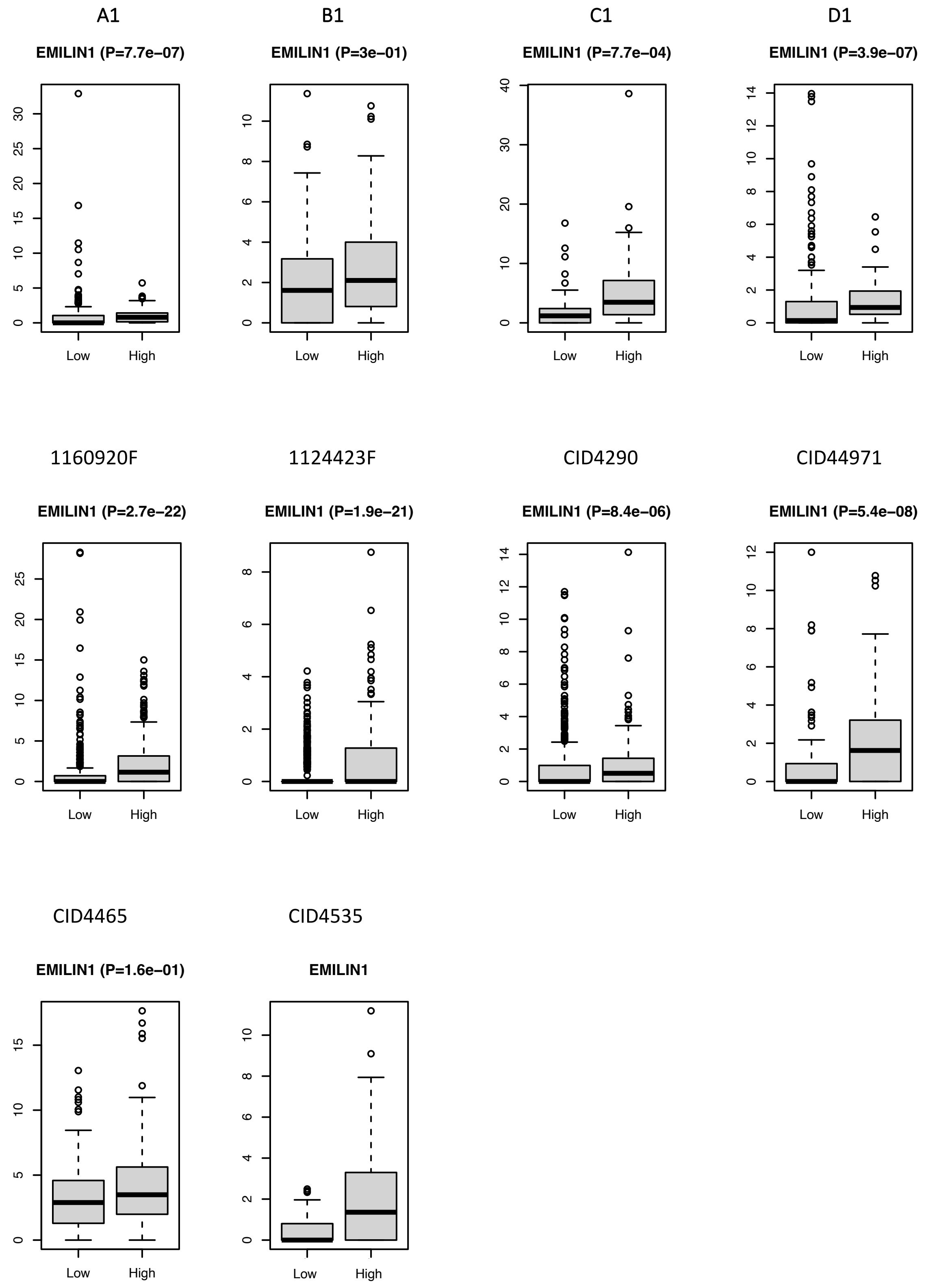
A1
EMILIN1 (P=7.7e−07)
B1
EMILIN1 (P=3e−01)
C1
EMILIN1 (P=7.7e−04)
D1
EMILIN1 (P=3.9e−07)
1160920F
EMILIN1 (P=2.7e−22)
1124423F
EMILIN1 (P=1.9e−21)
CID4290
EMILIN1 (P=8.4e−06)
CID44971
EMILIN1 (P=5.4e−08)
CID4465
EMILIN1 (P=1.6e−01)
CID4535
EMILIN1
Low
High

Figure S14
